# Microplate Format Protein Nanopatterning for High-Throughput Screening of Cellular Microenvironments

**DOI:** 10.1101/2023.11.19.567703

**Authors:** Ali Shahrokhtash, Malthe von Tangen Sivertsen, Sara Hvidbjerg Laursen, Duncan S. Sutherland

## Abstract

An advanced protein nanopatterned cell culture platform is engineered to emulate the extracellular matrix’s complexity, enabling precise nanoscale biomolecule copatterning to mimic environments analogous to native tissue for cellular assays. Nanopatterns fabricated through sparse colloidal lithography, with 100 nm to 800 nm features in separate wells, are seamlessly integrated into standard microplate formats (96-well/384-well). Robust patterns are built from fully PEGylated, passivated thin glass coverslips optimized for minimal nonspecific interactions. Biotin-avidin binding and click chemistry to ensure the accurate and robust localization of bioligands. The transparent, metal-free substrates are free of topographical interference, rendering them ideal for diverse fluorescence microscopy techniques encompassing single-molecule TIRFM and extensive high-throughput imaging. The structural stability of these nanopatterns persists beyond a year in storage and long-term in cell culture conditions, endorsing their application for prolonged experimental studies and potential for widespread academic and industrial use. The platform has been demonstrated for nanopatterning an array of biomolecules, from small molecules to proteins, DNA, and extracellular matrix components, instrumental for studying cell signaling. Experiments with C2C12 cells demonstrated the exceptional specificity of the nanopatterned microplates, with nonspecific adhesion remaining below 2% and the platform’s ability to elicit size-dependent cellular reactions when interfaced with nanopatterned fibronectin.

## 1. Introduction

The microenvironment of a cell and the interactions with the extracellular matrix (ECM), soluble factors, and contacts with neighboring cells define a set of extrinsic signals that, when combined with intrinsic cellular signals, determines cellular phenotypes.[1]

*In vitro* cell culture provides model systems and cellular assays by culturing cells taken out of their natural environments, typically in polymeric culture wells. These types of *in vitro* cell culture systems underpin almost all human efforts to understand and exploit the molecular biology of life in health sciences and medical technologies.

A key benefit of *in vitro* culture is the potential for reproducibility and the ease of access to modern analytical tools, such as high-resolution fluorescence microscopy.[2],[3]

While the typically polymeric culture wells do not themselves give specific signals to the cells, some of the microenvironmental signals can be provided by other cells in the culture, whether of the same or another cell type (*e.g.*, in co-cultures) or from soluble signaling molecules added to the cell culture media.[4]–[6]

In addition, cultured cells, over time, synthesize and release ECM proteins and soluble factors to condition their microenvironment over the time scale of hours to days. Together with proteins present in the media (often bovine serum containing), these can nonspecifically adsorb to the polymeric well surfaces.[7].

A widely used approach to provide more control of the microenvironment can be given by pre-adsorbing ECM proteins either composed of a single type of protein (*e.g.*, fibronectin) or complex mixtures (*e.g.*, collagens/Matrigel) to provide specific signals at the initial interaction of cells with the culture system (seconds to minutes) to provide simplified biochemical ECM mimics.[8]

However, providing appropriate cellular microenvironments to maintain or drive cells into phenotypes by mimicking particular *in vivo* environments remains a major challenge. Over the last decade, it has become clear that the physical properties of the microenvironment, such as stiffness, contribute to biomechanical signals activating mechanotransduction pathways, where polymeric materials are inappropriate to mimic most *in vivo* situations.[9],[10]

Efforts to generate more relevant biomechanical signals in culture systems with modified material properties for 2D- and 3D-culture systems based on soft protein-based or synthetic hydrogels and *via* spheroids/organoids are underway, giving advantages in complexity with the potential capability of better mimicking some types of *in vivo* microenvironments[11]–[13]. However, these approaches typically have significantly reduced analysis capabilities, *e.g.*, high-resolution imaging, high throughput, and automated processing.

Alternative approaches have suggested surface topography[14],[15] or surface biochemical micro- and nanopatterning[16]–[20] as routes to stimulate and control mechanotransduction. A concept being that *in vivo* cells and their microenvironment are organized at the molecular and nanometer scale, and synthetic materials that match those length scales can provide more relevant signals.

Topographic patterns have had an advantage in terms of scalability, as they can be integrated into the pre-existing processing used to produce polymer culture plates, and commercial arrays of nanotopographies are available and have been applied to the optimization of stem cell differentiation.[15],[21]

However, despite significant efforts, a mechanistic understanding of the role of nano and microtopographies is still lacking.

In contrast, surface-bound proteins can provide better understood and specific signals. Nanopatterning has emerged as a promising strategy to mimic cellular microenvironments and select and steer cellular adhesion complex formation[22]–[26] and mechanotransduction signaling[27] to generate specific cellular phenotypes.[25],[28]

Technologies for micropatterning proteins at the cellular length scales *in situ* in culture wells exist but suffer from slow write speeds and complex technology[29]. Nanoscale organization of proteins at the length scale of subcellular complexes is a powerful approach to control cellular adhesion and signaling at cellular adhesions, but to date, it has not been demonstrated in a platform compatible with high-throughput imaging (transparent, well-plate format with controlled protein immobilization and with a protein-rejecting underlay).

Here, we demonstrate a new robust technology for the formation of protein nanopatterns for copatterning of multiple proteins in a standard multiwell microplate format (*e.g.*, 96- or 384-well plates) for immobilization of ECM proteins and cellular receptors for cell culture experiments. The technology is based on a transparent, fully PEGylated, and metal-free substrate and can be applied with multiple protein and biomolecule immobilization strategies. This technology opens the way for the use of nanopatterning protein signals to define cellular phenotypes in a format compatible with high-throughput approaches. The practicality of this method is demonstrated through imaging by Total Internal Reflection Fluorescence

Microscopy (TIRFM), Confocal Laser Scanning Microscopy (CLSM), as well as high-throughput imaging of proof-of-concept cell experiments using C2C12 myoblasts.

## 2. Results & Discussion

Our goal was to develop a platform technology enabling the use of nanopatterns of proteins in a multiwell plate format. The key requirements were that the technology should allow the simple preparation of protein nanopatterns within sterile cell culture facilities in one-step processes and that the platform should be transparent and fully applicable for high throughput fluorescence and brightfield imaging in the same way as standard cell culture well plates. Our approach has been to fabricate multiwell culture plates with nanopatterned biofunctionalization groups on an anti-fouling surface that can be transported and stored long-term, then functionalized as needed with selected proteins and used in cell culture studies.

### 2.1 Transparent and Topography-Free Nanopatterning Fabrication for Multiwell Plate Formats

To fabricate nanopatterns on the surface, we use sparse colloidal lithography[30] (SCL) to pattern a mask material. The mask material (Cr) is then used to direct the assembly of patterns of two functionalized polymeric brushes *via* a lift-off and backfilling process. The polymeric brushes are selected to provide ultralow nonspecific protein binding and include click chemistry groups for site-specific immobilization of proteins and biomolecules of interest.

Sparse colloidal lithography as an approach has the advantage of not requiring a spin-coating step, as required for other current protein nanopatterning approaches, for example, in electron beam lithography[18],[31], UV-photolithography[17],[29], nanoimprint lithography[16],[32], block copolymer micelle lithography[20],[33],[34] and hole-mask colloidal lithography[26],[35],[36] which simplifies the process and makes scaling up to a well-plate format (78 mm x 111 mm) practical.

In brief, sulfate latex (polystyrene) nanoparticles, with a set of sizes from 80 nm to 800 nm and low polydispersity (<10%), were used to form sparse colloidal monolayers on different thin glass substrates *via* electrostatic self-assembly.

Prior to the electrostatic adsorption of the particles to the surface, we thoroughly cleaned the substrates and precoated them with 3 self-assembled monolayers of organic polyelectrolytes to achieve a thin adhesive layer with a stable net positive surface charge. The negatively charged sulfate-terminated latex nanoparticles are then allowed to adsorb to the surface to form dispersed colloidal monolayers. A two-dimensional random sequential adsorption[37] (2D-RSA) model can describe the particle adsorption process. When the particle coverage approaches the jamming limit, *i.e.*, when the available space for other particles to absorb is nearly exhausted, the adsorbed nanoparticles exhibit short-range ordering with a characteristic spacing resulting from the electrostatic repulsion between particles.[30],[38] However, the arrays of particles do not display the long-range order characteristic of crystallinity. Allowing the assembly to reach saturation levels, where the short-range order is well-defined, is recommended for most applications. Depending on the nanoparticle size and concentration, the process takes from 1 minute to 1 hour. (Figure 1a)

**Figure 1.**
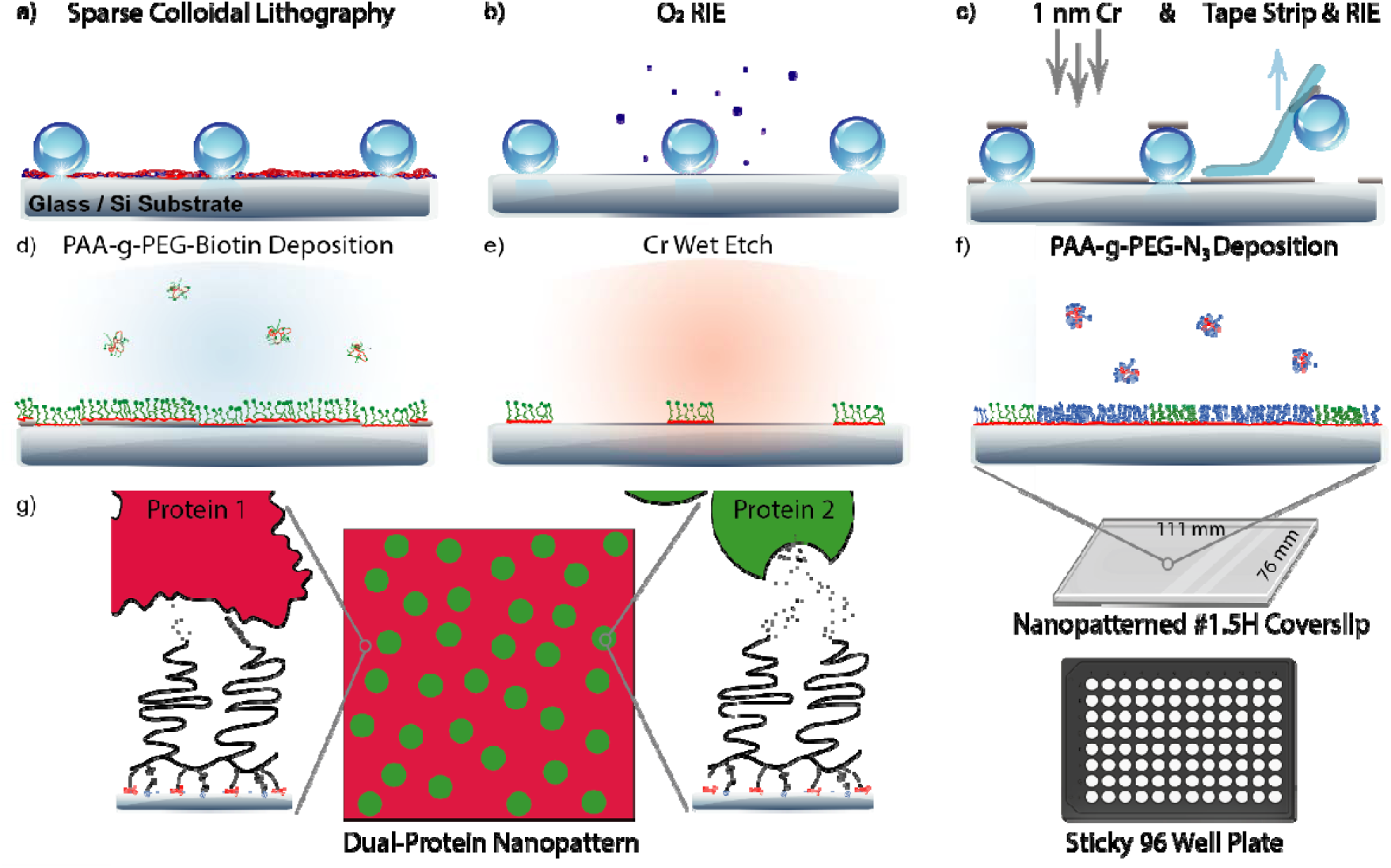
Fabrication steps for nanopatterning 2 ligands on a fully PEGylated topography-free background. a) After cleaning the substrate, polyelectrolyte layers are deposited to establish a net positive charge. A sparse monolayer of negatively charged polystyrene nanoparticles forms the mask for the following fabrication steps. b) A short O_2_ plasma RIE removes the organic polyelectrolyte layers, facilitating strong binding of the anti-fouling polymer PAA-g-PEG *via* NH_2_ and silane groups in the subsequent steps. c) A 1 nm Cr layer is deposited on the substrate, and the particles are removed by tape stripping, providing apertures directly onto the substrate. Another short O_2_ plasma RIE step removes the particle residues at this step. d) The anti-fouling random cograft polymer terminated with a Biotin-tag (PAA-g-PEG-Biotin (NH_2_, Si)) is introduced from an aqueous solution, forming a monolayer. e) A 2½ min Cr wet etching process removes the Cr layer and the PAA-g-PEG-Biotin bound to this layer, leaving nanoscale patches of PAA-g-PEG-Biotin corresponding in diameter to the size of the nanoparticles. f) Deposition of another biospecific PAA-g-PEG-N_3_ polymer on the surface allows for orthogonal binding of two bioligands on the surface while preserving the anti-fouling properties. After this stage, the multiwell-sized (111 mm x 76 mm x 0.17mm) nanopatterned thin glass substrate can be attached to a sticky-96-well plate for protein incubation. g) A schematic representing the top view of the patterned substrate after protein incubation. The inserts show the anti-fouling polymer’s covalent siloxane and electrostatic attachment to the glass. The biorthogonal tag is visualized to attach the biomolecule of interest with a DBCO-labeled protein (Protein 1) and a biotin-binding protein (Protein 2).

In the next step, O_2_ plasma reactive ion etching is briefly performed to remove the organic polyelectrolyte layer between the adsorbed particles of the colloidal monolayer mask. Following this, a 1 nm thin chromium (Cr) film is immediately deposited by physical vapor deposition. (Figure 1b-c and S1).

The Cr thin film covers the top of the particles and the gap between the adsorbed particles. Consequently, the particles act as a mask, preventing the area beneath them from being coated with Cr. Upon removal of these particles, nano-sized apertures with dimensions equivalent to the particles are formed directly on the glass substrate. Another brief O_2_ plasma reactive ion etching process removes the residual polyelectrolyte layer and potential debris from the nanoparticle, ensuring that the nano-apertures are clean and prepared to support the aqueous deposition of a random graft copolymer of amine, silane, and polyethylene glycol (PEG)[39],[40] onto the surface, which is terminated with a chosen biospecific tag, *e.g.*, biotin. The deposited polymer forms a firm bond with the surface *via* electrostatic attraction, facilitated by the positively charged amines and covalent interaction through the silane linkers. This dual-attachment approach ensures strong adhesion and durability.[39],[41]

As a result, PEG brushes are formed across the entire surface, both on the Cr film between the particles and inside the nano-apertures directly attached to the glass substrate.

The surface exhibits anti-fouling properties at this stage, meaning it resists nonspecific protein binding. However, it permits specific attachment of proteins to the tag (*e.g.*, avi-tagged proteins to biotin or click chemistry groups, *e.g.*, surface-bound azide to react with the DBCO functionalized proteins covalently).

The Cr thin film can be removed by wet chemical etching in a lift-off step, leaving the polymer brushes bound to the SiO_2_/glass substrate intact (Figure 1e) but removing the remaining PEG brushes. A clean SiO_2_/glass surface is revealed in the regions under the Cr layer. The PEG brushes, attached electrostatically and covalently to the glass surface, form nanoscale patches of brush polymer corresponding in diameter to the size of the nanoparticles used. At this stage, the nanopatterned section of the surface is carpeted with PEG brushes. In contrast, the rest of the surface is receptive to binding with an additional orthogonal biospecific tag-terminated PEG brush (Figure 1f). This arrangement of orthogonal biospecific tags facilitates the nanopatterning of two distinct ligands - one within the former locations of the nanoparticles and another in the spatially defined regions between the particles. If the specific experimental conditions do not necessitate the arrangement of molecules between nanopatterned regions, utilizing anti-fouling polymers such as PAA-g-PMOXA can be a viable alternative at this stage, given their notable stability.[41]–[43] As a result, we obtain a nanopatterned surface that is fully transparent, metal-free, and free of topographical features while exhibiting anti-fouling properties. A representation of the final product post-incubation with tagged bioligands is shown in Figure 1g. These features are critical for managing specific interactions at the engineered biointerface, especially when utilizing complex media such as serum-supplemented cell cultures, which have the potential to adhere nonspecifically to the surface and replace molecules bound nonspecifically – a common attachment method in other nanopatterning strategies.[19],[23],[26],[44]–[46]

Two primary factors influence the quality of the nanopatterned surface. The first is the adsorption of the nanoparticles. Once the colloidal nanoparticles are deposited from the aqueous solution, the surface must be dried for subsequent patterning steps. During this process, capillary forces can induce aggregation of the particles, particularly those larger than 200 nm (Figure S2). To counteract this, two simple strategies can be implemented: incorporating solvents such as ethanol into the solution to lower surface tension or heating the substrate in an autoclave or a microwave oven beyond the glass transition temperature (T_g_) of polystyrene, thereby expanding the contact area between the particles and surface, thus reducing the impact of capillary forces In our experiments, we found that heating above the T_g_ proved sufficient, although larger particle sizes might necessitate employing both strategies simultaneously. The second factor impacting the nanopatterning quality is the wet etching duration of the Cr layer. A continuous Cr layer is required for the patterning process to prevent the leakage of the first deposited anti-fouling polymer in the gaps between the nanopatterned regions. However, an extended Cr etching period would be required if the deposited Cr film is overly thick. Long etching times have the potential to allow oxidative damage to the already formed PEG brushes, which could significantly deteriorate that part of the surface’s anti-fouling characteristics.

Similarly, the biospecific tags on the PEG brushes may also be compromised. Multiple tags are possible to use, but we recommend that the most robust tag, in this case, biotin, be best placed first within the nanopatterned regions exposed to the Cr etchant. Therefore, using the thinnest possible layer and the shortest possible Cr wet etching time is optimal.

We validated the removal of Cr layer using visual inspection and X-ray photoelectron spectroscopy (XPS) (Figure S3-4). Similarly, the anti-fouling properties and functionality of the first biospecific tag following the Cr etching were verified to tolerate 2½ minutes of Cr etching. This was demonstrated using Surface Plasmon Resonance (SPR) on SiO_2_-covered SPR chips, which were prepared identically to the patterned glass surfaces. These experiments highlight the capability of the polymers exposed to the Cr etchant solution to prevent the nonspecific adhesion of over 98% of the proteins in the solution compared to the bare SiOLJ surfaces while still maintaining the specific biotin-avidin interaction (Figure S5-S7).

We prepared patterns on glass substrates of different sizes, from 25 mm diameter glass to whole multiwell plate size (78 mm x 111 mm). Following this, the multiwell format nanopatterned substrates were glued to custom-made sticky-96-well plates (Figure 1f, S8) or commercially available 384-well plates using silicone as the adhesive. As an alternative option, to encompass various nanopatterning dimensions on one substrate, the glass substrate prepared during the fabrication step ’C’ can be segmented into smaller units and adhered to the same multiwell plate to give 2, 4, 8, etc., different patterns in a single well plate.

The transparent substrates formatted for well-plate applications were effectively utilized in subsequent steps for widefield fluorescence microscopy, CLSM, TIRFM, and high throughput widefield imaging. Across the extensive >85 cm² substrate, we noted a consistent pattern quality, absence of topographical features (Figure S9), and well-maintained anti-fouling properties. Throughout the tested 10-day incubation period at 37°C, no leakage between the custom-made 96-well plate wells was detected, substantiating this platform’s suitability for extended *in vitro* cell experiments.

### 2.2 Nanopattern Formation Across Size Ranges

We demonstrated the ability of our patterning technique to produce nanopatterns of different sizes. We employed Scanning Electron Microscopy (SEM) to evaluate the establishment of the sparse colloidal monolayer alongside its organization and coverage across the multiwell plate-sized coverslips used in the fabrication steps. As previously illustrated by Hanarp et al. [30],[38], a sparse monolayer of particles was assembled, exhibiting short-range order across all sizes studied. Representative images for the size range (80 nm to 800 nm) are shown in Figure 2a.

**Figure 2.**
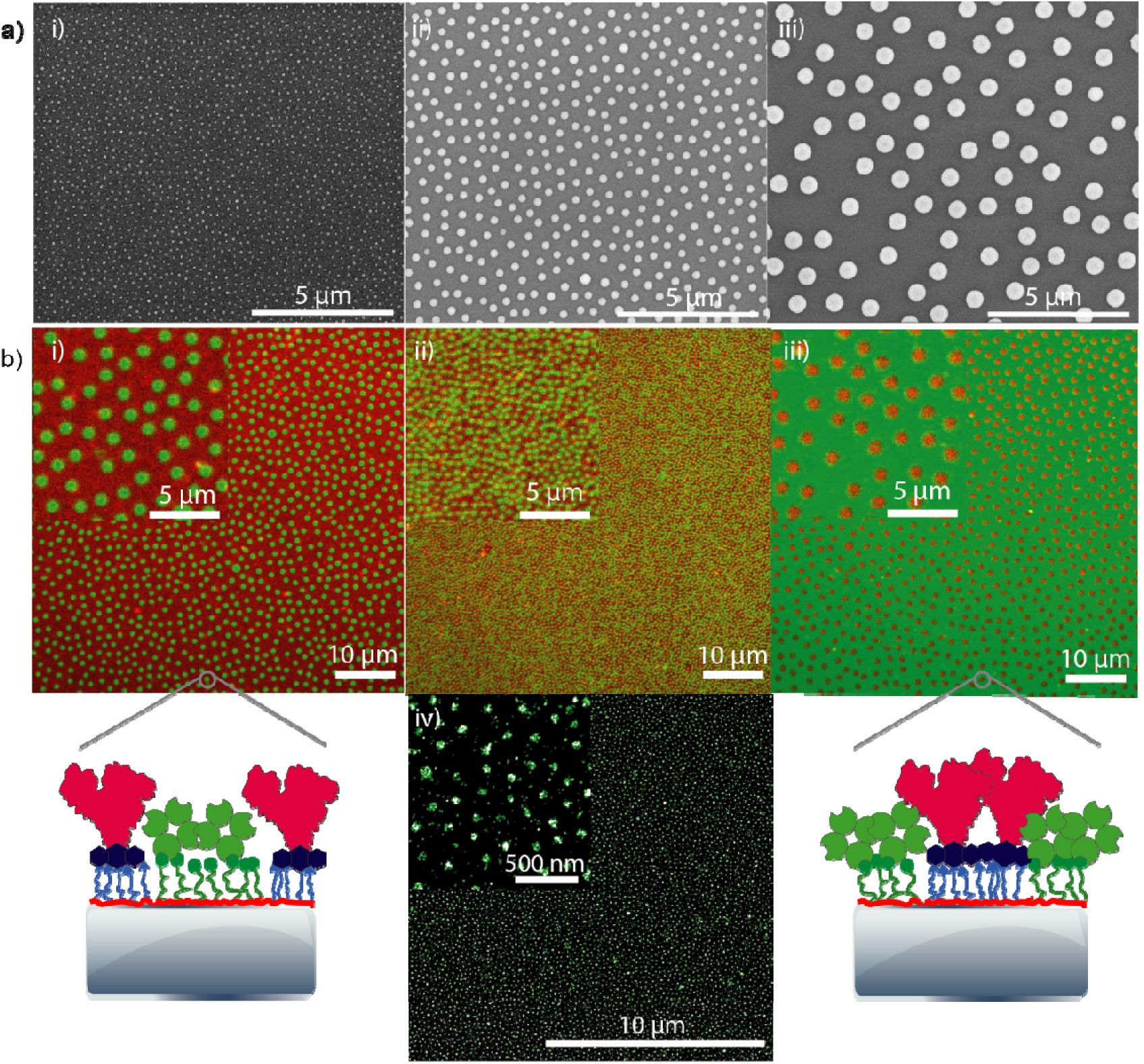
Nanoparticle assembly and protein pattern formation across several size ranges. a) SEM images of particles assembled on the surface. Features: i) 80 nm ii) 300 nm iii) 800 nm particles. b) Verification of protein nanopattern formation through confocal laser scanning fluorescence microscopy (CLSM) and DNA-PAINT super-resolution microscopy i) CLSM image of 800 nm protein patterns ii) 300 nm and iii) 800 nm protein patterns with reversed-tags on the surface. iv) DNA-PAINT super-resolution image of 80 nm streptavidin. The top left inserts are a magnification of the same surfaces. Green indicates Streptavadin, and red DBCO-BSA, used to demonstrate the patterning system.

Next, we validated the generation of the nanopatterns by using fluorescently labeled. streptavidin (SA) and Dibenzocyclooctyne (DBCO) -functionalized fluorescent bovine serum albumin (DBCO-BSA) to site-specifically assemble at a PAA-g-PEG-Biotin/PAA-g-PEG-N_3_ patterned surface. For the system to function as intended, we anticipate that SA will bind to the PAA-g-PEG-Biotin brushes within the nanopatterned region. At the same time, DBCO-BSA covalently attaches *via* a biorthogonal strain-promoted azide-alkyne cycloaddition[47] (SPAAC) to the terminal azide groups on the PAA-g-PEG-N_3_ brushes [48] on the surface in the region between the nanopatterns. Furthermore, it is expected that the complete formation of the protein layers within the predetermined areas would occur at different time scales. This becomes evident when comparing the second-order reaction rate of avidin-biotin (10^7^ M^-1^ s^-1^)[49] to that of DBCO-N_3_ SPAAC (0.1 M^-1^ s^-1^). [50] The SA layer assembly is predicted to be complete within minutes following incubation at the concentration range used (25 μg/mL ≈ 0.5 μM), in stark contrast to the DBCO-labeled protein incubation, which necessitates elevated concentrations and temperature conditions for optimal results.

In our findings, an overnight incubation of DBCO-labeled proteins at an approximate concentration of 1 μM (≈ 100 μg/mL for DBCO-BSA) at 37°C culminates in surface saturation. For scenarios where quicker reaction rates are desired, the substitution of DBCO with alkyne tags could be a viable strategy. This adjustment allows for a covalent interaction with the surface’s N_3_ group through Copper(I)-catalyzed Azide-Alkyne Cycloaddition (CuAAC), with a 100-fold higher second-order reaction of 10 M^-1^ s^-1^.[51]

Nonetheless, we chose to utilize the Cu-free SPAAC reaction given its straightforward approach and a diminished risk of copper toxicity affecting the subsequent *in vitro* experiments.[52]

Considering the aforementioned factors during surface preparation proves to be crucial. Figure 2b i-ii illustrates that the protein nanopatterns have successfully assembled within the predefined regions. CLSM images were used for samples with patterns larger than the diffraction limit. This is further supported by a Pearson correlation coefficient of -0.5, indicating a robust anti-correlation between the two fluorescence channels shown in the line profiles representing the fluorescence signal intensities in Figure S10.

We also demonstrated that it is possible to swap the pattern between the nanopatterned domains and the intervening background regions (Figure 2b iii). This was done by altering the biospecific tags deposition sequence: PAA-g-PEG-Biotin and PAA-g-PEG-N_3_. This adjustment in the procedure seeks to provide nuanced control over the spatial distribution of the molecules without relabeling the biomolecules of interest with biotin and DBCO. While this "tag-swapping" patterning is technically possible, we observed that the biotin molecules maintained greater stability throughout the nanofabrication process. Consequently, we recommend altering the tags on the biomolecules instead of on the surface, whenever feasible, for optimal results.

Having confirmed the formation of nanopatterns above the diffraction limit, we utilized DNA-PAINT[53],[54] super-resolution microscopy to verify the generation of sub-diffraction limited nanopatterns. For this, a short DNA docking strand was conjugated to SA. By localizing the transient binding events of a complementary DNA oligo with a fluorophore across 10,000 frames, we reconstructed a super-resolution image of the 80 nm wide SA nanopatterns (Figure 2b iv), confirming the formation of sub-diffraction nanopatterns.

Indeed, while the current experiments utilized nanoparticles with diameters ranging from 80 nm to 800 nm, the smallest feature size achievable with this nanopatterning strategy can be much smaller. This is mainly constrained by the size of the anti-fouling polymer used. For instance, a similar polymer’s footprint has been reported to be approximately 70 nm^2^. [32] This would suggest that the smallest possible circular nanostructure achieved with this approach could have a minimum diameter as low as 10 nm. This indicates the potential to use this technique to construct nanopatterns with single molecules, further enhancing the method’s potential application. However, given that commercially procured colloidal particles below 100 nm typically exhibit high polydispersity, sparse colloidal lithography might not be the best fabrication technique for creating such small structures. An alternative approach could involve minimizing the degree of Biotin molecule functionalization on the PEG brush. This would allow the 80 nm structures currently being utilized to capture, on average, one molecule.

Having established the successful formation of protein nanopatterns across the utilized size range, we investigated whether particle density and spacing can be modulated. This is of significance, as some specific biological processes exhibit sensitivity to the global density of the ligands.[22],[55] Allowing the nanoparticles to approach the jamming limit on the surface by incubating at comparatively high concentrations for extended periods enables maximal nanostructure density on the surface. Conversely, prematurely abrupting the process before reaching the jamming limit, through either reduced particle deposition duration or lower particle concentrations, decreases surface coverage. However, under these circumstances, the short-range order of structures may be lost due to the stochastic nature of particle arrival on the surface (Figure S11). Nevertheless, this approach can facilitate surface coverages below 10%.[30] Similarly, Andersen et al. showed that utilizing sparse colloidal lithography by employing angled deposition facilitates the creation of ring and crescent-like structures instead of circular disks, thereby modulating nanopattern coverage from 8% to over 30%. [44]

To alter the overall nanostructure coverage while preserving the short-range order and the characteristic inter-structure spacing, modulation of the net surface charge and ionic concentration during the particle solution phase could prove beneficial.

Figure 3 exemplifies the impact of altering the concentration of the last polyelectrolyte layer deposited onto the surface prior to the adsorption of the nanoparticles. By increasing the concentration of the positively charged Poly(diallyldimethylammonium chloride) (PDDA) polyelectrolyte layer while keeping all other parameters constant, the coverage of the 210 nm nanoparticles was modulated from 12.8±0.2% to 21.7±0.4%. Despite this change, the short-range order among the particles was maintained.

**Figure 3.**
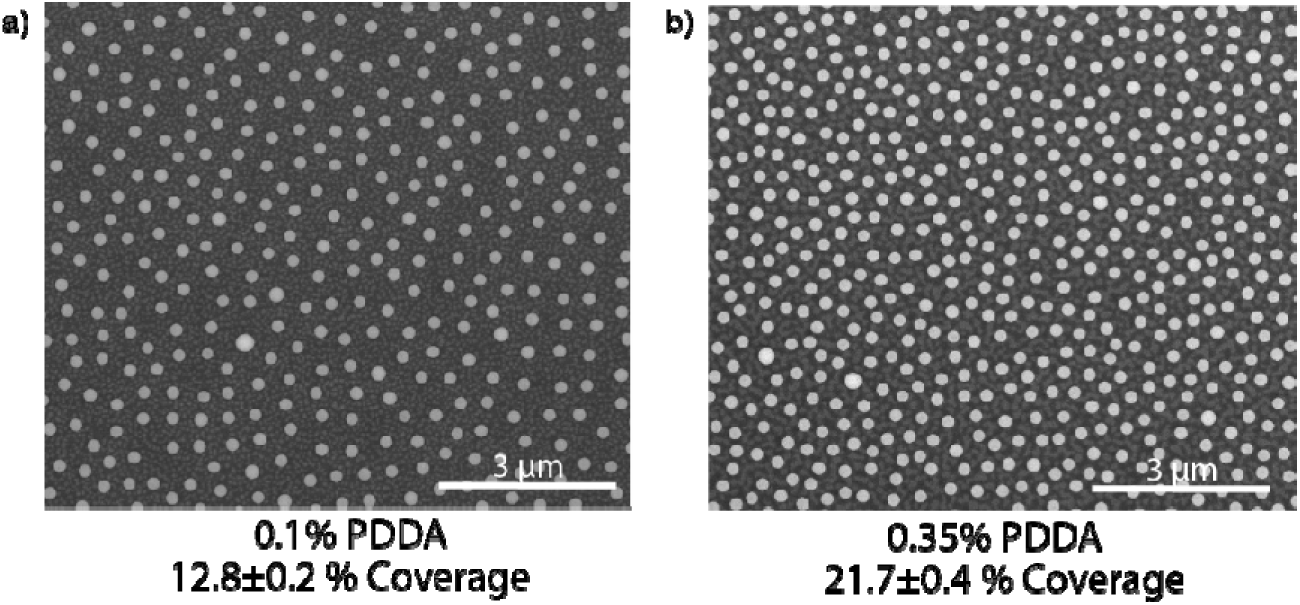
Altering the surface charge tunes the distance between the adsorbed nanoparticles. Representative SEM images of 0.2 wt.% 210 nm polystyrene nanoparticles deposited for 10 minutes on different concentrations of PDDA, the final polyelectrolyte layer applied to the surface. By tuning the PDDA concentration, the surface coverage of the adsorbed nanoparticles is modulated from 12.8% ±0.2% (a) to 21.7±0.4% (b). In contrast, the inherent pattern and short-range order - governed by the random sequential adsorption model depicting the stochastic arrival of particles on the surface - remain preserved.

While the increase in PDDA concentration significantly increased the surface coverage in this instance, there seems to be an upper limit to this effect. If higher surface coverages are desired, another strategy is to counteract the Coulombic repulsion between the negatively charged polystyrene nanoparticles adsorbed on the surface and those arriving at the surface, allowing them to land closer to each other. Hanarp et al. demonstrated that adding μM monovalent ions to the particle solution can further increase the particle density on the surface. Similarly, by shrinking the nanoparticles using reactive ion etching, even lower coverages can be achieved while maintaining the short-range order. [30],[38]

### 2.3 Biomolecule Nanopatterning Diversity and Stability

Having verified the low nonspecific binding and specifically patterned chemistry across several size ranges, we explored the method’s adaptability for patterning various biomolecules. Additionally, we examined its compatibility with different fluorescence-based imaging techniques and assessed the stability of the formed patterns.

A prevalent category of proteins used in *in vitro* assays are high molecular weight ECM proteins, including fibronectin (FN), which is typically nonspecifically adsorbed at a surface and often involves partial denaturation.[56] Nanopatterning of FN has previously been demonstrated to steer epidermal stem cell differentiation.[25] In Figure 4a, we demonstrate the efficient patterning of FN on a PEG brush as visualized by an inverted widefield epifluorescence microscope. Biotinylated FN, available commercially or prepared through random biotinylation of lysines using carbodiimide crosslinkers such as N-hydroxysuccinimide (NHS) esters, rapidly binds to the SA molecules bound to the PAA-g-PEG-Biotin brushes patterned on the surface. The tetrameric SA molecules create a linkage between the nanopatterned Biotin tags on the surface and the Biotin tags present on the Biotinylated FN molecule, facilitating FN adherence within the patterns atop the SA molecules. We also confirmed the formation of the FN layer using SPR, as shown in Figure S12.

**Figure 4.**
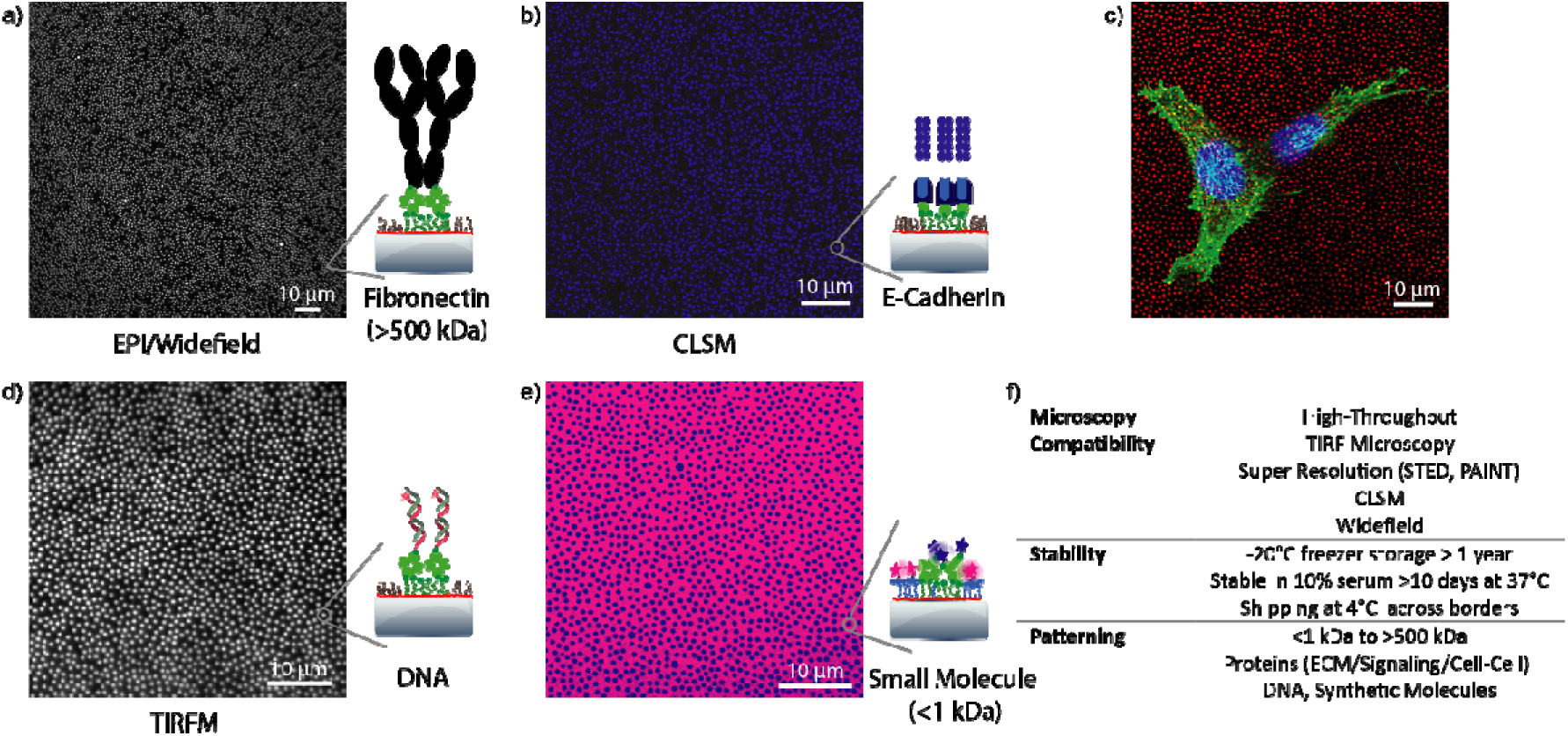
The nanopatterning methodology facilitates the patterning a broad spectrum of biomolecules and is compatible with all fluorescence microscopy techniques. a) Epi/Widefield inverted fluorescence microscope image of 600 nm structures featuring patterns formed by fluorescently labeled biotinylated-fibronectin (M*_w_* > 500 kDa), with streptavidin acting as a linker protein between the patterned PAA-g-PEG-Biotin molecules on the surface and the biotinylated-fibronectin. b) CLSM image illustrating the orientation of the cell-cell junction protein E-Cadherin:Fc on 500 nm structures, mediated through biotinylated Protein A and streptavidin as linker proteins. d) 3T3 fibroblasts adhering to fibronectin 600 nanopatterns visualized by immunostaining. Red: fibronectin, Green: F-Actin, Blue: Nucleus d) TIRFM image capturing fluorescently labeled DNA molecules on 600 nm nanopatterns, facilitated through a streptavidin linker and a terminal biotin tag present on the DNA molecule. e) CLSM image providing a proof-of-concept for the patterning of molecules below 1 kDa in size. Pink denotes DBCO-5’FAM, and blue represents Cy3-Biotin. f) Summary of the system’s compatibility and stability aspects. The surfaces in figures a,d and b utilize PAA-g-PMOXA as the anti-fouling background polymer.

Figure 4c illustrates the interaction of the 3T3 fibroblast cell lines with the FN nanopatterns in a cell culture environment after 24 hours, validating the functionality and stability of the FN-nanopatterns. We also found that these nanopatterns retain their pattern and function for the 10-day tested period in cell culture media supplemented with 10% serum at 37°C, allowing long-term cell culture experiments on the nanopatterned substrates (Figure S13).

To further assess the chemical stability of the patterned surfaces, we prepared and shipped fully assembled 96-well plates with nanopatterned coverslips of various sizes to an overseas collaborator. Throughout the transit, the plates were maintained at 4°C, followed by a storage period of up to 3 months in a -20°C freezer, housed in sealed bags filled with N_2_ to mitigate potential oxidative damage to the PEG brushes.[57] Remarkably, we observed no discernible degradation in the qualitative attributes of the outcomes derived from these nanopatterned substrates.

Furthermore, we stored a patterned coverslip under analogous conditions for over a year (561 days), preserving its excellent anti-fouling characteristics while still allowing the specific interaction of tagged biomolecules within the predefined regions. (Figure S14)

Another category of widely used molecules for *in vitro* assays, especially in the study of cell-cell interactions, are cell-cell junction proteins.[58] The CLSM image in Figure 4b illustrates the nanopatterning of the cell-cell junction protein E-cadherin. To orient E-cadherin in a physiologically relevant orientation on the surface, we employed a chimeric construct comprising monomeric streptavidin (mSA) and staphylococcus aureus Protein A (PrA)[59], a Fc binding protein (Figure S15, S16).

The mSA is arranged on the nanopatterned PEG-Biotin brushes on the surface. At the same time, the PrA moiety allows the binding of an Fc tag genetically engineered onto the E-cadherin protein with high affinity. This arrangement ensures a biologically relevant orientation of the E-cadherin protein on the surface, which is essential for modulating the cellular behavior at the nanoscale.[60]

Although the mSA-PrA conjugates expedite the surface preparation process, a more straightforward combination of sequentially incubating the surface with SA and biotinylated PrA can also orient Fc-tagged proteins. This has been demonstrated in our prior work for the orientation of ICAM-1 on nanopatterned surfaces.[44] We confirmed the immobilization using SPR in Figure S17.

The versatility of our nanopatterning approach is not limited to proteins. Another subset of biomolecules of interest is DNA. Synthetic DNA molecules have gained significant interest in biological applications, including biosensing and therapeutics. We further extended our validation to these molecules by binding fluorophore-conjugated DNA, functionalized with a terminal biotin tag, and SA as the linker between the surface and the biotinylated DNA. These structures were observable using TIRFM, which offers the potential to study the interaction of DNA molecules with their molecular targets or cells in 100-200 nm thick axial sections proximate to the nanopatterned surface. Compatibility with this imaging modality further suggests the potential of the patterning technique for application in single-molecule fluorescence studies.

Last, we turned our attention to the patterning of small synthetic molecules, typically below 1 kDa. These molecules, often holding therapeutic relevance[61], represent another class of biologically pertinent ligands characterized by their smaller dimensions and molecular weights compared to proteins. To confirm whether these small molecules would bind selectively to the spatially defined regions, we tested the binding of DBCO and biotin-functionalized fluorophores with molecular weights under 1 kDa to the surface. The successful execution of this is demonstrated in Figure 4e. A summary of the versatility, stability, and compatibility of the nanopatterning system with different fluorescent imaging techniques is given in Figure 4f.

### 2.4 Myoblasts Show Differential Adhesion Characteristics on Nanopatterned Fibronectin

Next, we prepared a 96-well plate with 3 different nanopatterned sizes featuring three distinct nanopatterned scales with a twofold order of magnitude variation in each patterned region area: 80 nm (5⋅10^-3^ μm^2^), 200 nm (4⋅10^-2^ μm^2^) and 800 nm (5⋅10^-1^ μm^2^). A non-patterned homogeneous surface covered with PEG-Biotin, resembling the nanopatterned regions on the other substrates, was also assembled to form a complete 96-well plate (Figure 5e).

**Figure 5.**
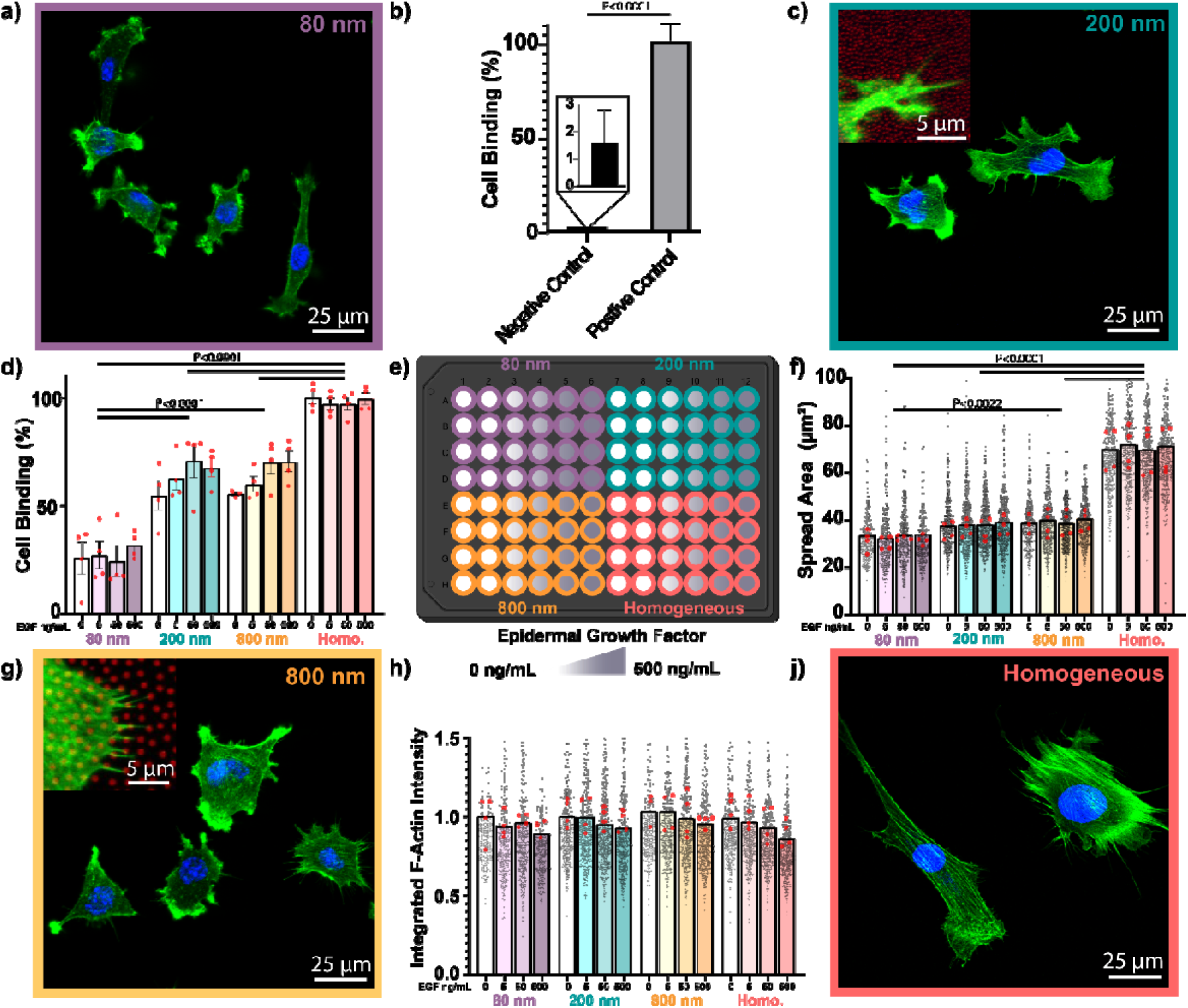
Myoblast cell adhesion and interaction with nanopatterned fibronectin (FN) across a 2 orders of magnitude area difference. a,c,g,j) Fluorescence image of C2C12 myoblasts spreading on different wells, representing different FN nanopattern sizes. The insert represents a zoom-in of the cellular interaction with the stained FN nanopatterns (red). Blue represents the nucleus, and green is F-Actin. b) Cell binding to Streptavadin negative control on homogeneous and nanopatterned substrates compared to cell binding to homogeneous FN substrate. d) Cell binding and seeding efficiency across size ranges and epidermal growth factor (EGF) concentrations. e) Overview of the plate with different nanopattern sizes and EGF concentrations. f) Quantification of the spread area. h) Quantification of integrated F-actin intensity normalized to 0 ng/mL EGF of the same pattern size. Red dots represent the mean value of each technical repeat (n=4), and gray dots represent the individual observations. The bars represent the median with standard deviation as the error bars. P values are calculated based on one-way ANOVA for d,f comparing the groups of the same pattern size, and Student’s t-test for b.

The plate was then covered with SA and biotinylated FN to assemble FN nanopatterns. As a negative control, a series of wells across all size ranges were covered only with SA, which lacks the cell adhesive motifs in FN.

As a proof-of-principle screening experiment, we cultured C2C12 mouse myoblast cells in the presence of 0 to 500 ng/mL epidermal growth factor (EGF). Post-culture, the cells were fixed, and the nuclei and F-actin were stained for imaging of the whole plate with a high-throughput fluorescence microscope, taking less than 1 hour to image the whole plate. The acquired data were analyzed using CellProfiler[62], an open-source image analysis software specialized for cellular microscopy data analysis. Fluorescence thresholding and object size-based filtering were used to define the nucleus, from which a mask was expanded to define each cell’s spread area from the F-Actin staining, enhanced by a trained pixel-classification machine learning model in Ilastik.[63]

The analysis revealed that less than 2% of the cells were able to bind nonspecifically to the substrates across different size ranges (Figure 5b) and EGF concentrations. These cells exhibited a highly circular morphology and limited spreading due to restricted interaction with the passivated substrates (Figure S18).

In the FN nanopatterned wells, the cells displayed a size-dependent adhesive and spreading behavior. Significantly fewer cells could adhere to the smallest 80 nm (5⋅10^-3^ μm^2^) FN patterns compared to all other larger sizes (Figure 5d). The size-dependent adhesion was also evident when the homogeneous substrate was compared with 200 nm (4⋅10^-2^ μm^2^) and 800 nm (5⋅10^-1^ μm^2^) patterns. These nanopatterned substrates also showed a trend of increasing adhesion efficiency at higher EGF concentrations (50-500 ng/mL).

The differential spreading area (Figure 5f) is also evident in the representative fluorescence images (Figure 5 a,c,g,j). The adhesion onto the smallest nanopatterns leads to the smallest spread area, with a trend of increasing spread area as the FN nanoclusters’ dimensions increased, leading to the most spread cells on the homogeneously covered FN substrate. The cells on the nanopatterned wells showed intense F-actin signals from few adhesion points, while the cells on the homogeneous substrates showed many stress fibers instead.

The differences in the F-actin intensity across all sizes showed a correlation with EGF concentration. Increasing the EGF concentration led to downregulation of the integrated F-actin intensity trend (Figure 5h).

The ultra-low nonspecific cell binding, in contrast to the typical 30-40% adhesion to BSA-blocked surfaces[64],[65], a common blocking method, facilitates control over cell interactions within the defined nanopatterned regions. This fact, coupled with the high-throughput compatibility of this nanopatterning platform, allows the development of high-throughput screening models, mimicking the complex *in vivo* cellular environment at the nanoscale to study the cellular interactions in the presence of up to 2 bound ligands and soluble drug/signaling molecules. This model system is free of potential interactions or artifacts attributable to nanotopography, commonly observed in other protein nanopatterning systems. This absence significantly improves the reproducibility of results concerning protein clustering at the nanoscale.

## 3. Conclusion

In conclusion, this study presents a novel and robust technology for patterning proteins, DNA, and other bioligands beneficial for co-patterning various relevant biomolecules at sub-100 nm to microns within standard multiwell microplate formats. This applied nanofabrication allows for low-cost, robust nanopatterning on a transparent, fully PEGylated, and metal-free substrate that integrates seamlessly with high-throughput analytical methods and single-molecule imaging approaches.

The ease of production of nanopatterns across multiple sizes with controlled surface coverage, the versatility of classes of biomolecule immobilization, long-term stability in culture, and ultra-low nonspecific cellular interaction lays a promising groundwork for fostering more nuanced understandings of cellular adhesion and signaling dynamics at subcellular scales, as well as the steering of cellular phenotypes to better mimic *in vivo* types. The compatibility of the nanopatterned wellplates with high throughput imaging gives the potential for developing phenotypically improved *in vitro* cellular assays for use in early drug discovery and validation or nanotoxicity screening.

## Supporting Information

Supporting Information is available online.

## Acknowledgments

We extend our gratitude to Fiona Watt’s Group for their diligent testing of the substrates’ robustness post-transport. We also thank Bjarke Rolighed Jeppesen for expertly sputter-coating the SiO_2_ SPR chips and Janni Nielsen for invaluable guidance in size exclusion chromatography.

This work was funded by the Danish National Research Foundation (DNRF135) and the Open Discovery Innovation Network (ODIN) project CELPPLUS sponsored by the Novo Nordisk Foundation (NNF20SA0061466).

## Data availability statement

The data that support the findings of this study are available from the corresponding author upon reasonable request.

