## Supplementary Material for "Microplate Format Protein Nanopatterning for High-Throughput Screening of Cellular Microenvironments"

#### Table of Contents:

Experimental Methods
S1: SEM side view of the Substrate before Cr etching
S2: SEM top view of capillary forces’ effect on particle assembly
S3: XPS of a substrate before and after Cr etching
S4:AFM and fluorescence scan of patterning non-continuous Cr layer
S5: Effect of Cr etching on the PEG-Biotin layer for Biotin-Streptavidin interaction
S6: Effect of Cr etching on unspecific protein binding
S7: Example SPR sensorgram for unspecific binding of BSA
S8: Photo of the 96-well plates and transparency of the nanopatterned substrates
S9: AFM scans of the substrate during the nanofabrication steps
S10: Line scan and anti-correlation of protein binding in patterns vs background
S11: Fluorescence image of kinetically abrupted particle binding, showing low nanopattern coverage
S12: Biotinylated Fibronectin binding to PEG-Biotin through Streptavadin as linker
S13: Fluorescence image of Streptavidin nanopattern after 10 days in serum
S14: Fluorescence image of nanopattern formation on >1 year-old sample stored in a freezer
S15: SDS-PAGE and size-exclusion chromatogram for ProteinA-Monomeric Streptavadin conjugate
S16: SPR of monomeric Streptavadin Protein A conjugate binding to PEG-Biotin
S17: SPR of Step-by-step binding of proteins to PEG-Biotin through Streptavidin, Biotinylated Protein A, and Fc fused protein
S18: Cell circularity data for the proof-of-concept cell experiment
S19: Flowchart for polyelectrolyte deposition on multiwell covering coverslips
S20: Flowchart for nanoparticle assembly on multiwell covering coverslips

### Experimental Methods

#### PEG-brush Functionalization

Poly(acryl-amide)-g-(PMOXA, 1,6-hexanediamine, 3-aminopropyldimethylethoxysilane) (4425:116.2:161.3 Mr; 0.20:0.40:0.40) PAA-g-PMOXA (NH_2_,Si) and poly(acryl-amide)-g-(PEG-N3, 1,6-hexanediamine, 3-aminopropyldimethylethoxysilane) (3500:116.2:161.3 Mr; 0.15:0.425:0.425) PAA-g-PEG-N_3_ (NH_2_,Si) were synthesized and characterized by SuSoS, Switzerland. Each polymer can be designed with multiple surface linkers; the linkers are specified in parentheses. PAA-g-PEG-N_3_ polymer's terminal azide (N_3_) group was modified with the bifunctional at least 5 times excess of DBCO-PEG_4_-Biotin (Sigma-Aldrich) compared to the estimated number of N_3_-groups. The PAA-g-PEG-N_3_ polymer was dissolved at 1 mg/mL in 1 mM (4-(2-hydroxyethyl)-1-piperazineethanesulfonic acid) (HEPES) (Sigma-Aldrich) buffer adjusted to pH 7.4 and was reacted overnight at room temperature with shaking in the dark. To remove the excess DBCO modifier, the polymer was filtered five times in an Amicon centrifugal filter with 30 kDa cut-off.
The modification's success was then confirmed by proxy in an SPR experiment.

#### Nanopatterning of Substrates

Coverslips (Ø25mm or 111mm x 77 mm - #1.5H) were cleaned by ultrasonication in acetone and isopropanol alcohol dried under a stream of N_2._ The substrates were treated with a 3-minute-long oxygen plasma reactive ion etching (RIE) (Vision 300 MK II, Advanced Vacuum) with a radio frequency generator power of 100 W, 100 SCCM O_2_ flow at 25 mTorr pressure before following a modified protocol ^[1]^ for sparse colloidal lithography (SCL) to assemble the nanoparticles on the surface. Three self-assembled monolayers (SAMs) of organic polyelectrolytes were deposited on the substrates to achieve a net positive surface charge. and First Polyethyleneiminee – branched M_w_ 25000 (Sigma-Aldrich) at 2 wt. %, followed by Poly(sodium 4-styrenesulfonate) M_w_ 70000 (Sigma-Aldrich) at 2 wt. % and finally Poly(diallyldimethylammonium chloride) M_w_ 200000-350000 (Sigma-Aldrich) at a concentration range of 0.01 wt.% to 0.5 wt. % depending on the desired surface coverage. Each solution was incubated for 30 seconds on the surface, followed by rinsing with DI H_2_O for 30 seconds and drying under a stream of N_2_. For the multi-well format coverslips, an ultrasonication rinsing and washing step was added to each step to ensure the removal of loosely bound polyelectrolytes (Figure S19). 80 nm to 800 nm negatively charged sulfate latex beads (ThermoScientific) were deposited at 0.1-0.2 wt. % (30 sec to 45 min) to form a sparse monolayer of nanoparticles (Figure S20). The unbound particles were removed by rinsing with DI H_2_O. Particles smaller than 200 nm were dried directly under a stream of N_2_. Ø25 substrates with particles larger than 200 nm were dropped in boiling water for 1 minute to increase the adhesion and then dried under a stream of N_2_. The multiplate-sized coverslips were heated in a 100 mL DI H_2_O bath for 10 minutes in a microwave oven (900 W) before drying under a stream of N_2_.

The deposited polyelectrolyte SAMs were removed with 1 minute of O_2_ RIE (50W, 100 SCCM O_2_, 25 mTorr) to use the deposited nanoparticles to fabricate the nanopatterns. E-beam thermal physical vapor deposition (PVD), Cryofox Explorer 500 (Polyteknik, Denmark) with a base pressure < 10^-6^ mTorr was used to deposit a 1 nm thin Chromium metallic mask from 78 cm at a rate of 0.08 Å/s with 5 RPM rotation. The nanoparticles were removed by tape stripping. 1 minute O_2_ RIE (35 W, 40 SCCM, 50 mTorr) removed polyelectrolyte SAM leftovers and nanoparticle residues under the removed particles.

A droplet of 0.1 mg/mL PAA-g-PEG-Biotin (100%) in a 1 mM HEPES buffer at pH 7.4 was placed on Parafilm. The samples were then placed on the droplet and incubated for 30 minutes with a subsequent 30 seconds of rinsing with DI H_2_O and drying under a stream of N_2_ gun. Subsequently, a 0.2 μm filtered ceric ammonium nitrate-based Chromium etchant solution (Sigma-Aldrich - 651826) was used for 2½ minutes to remove the chromium and chromium oxide layer. Samples were rinsed with DI H_2_O and dried under a stream of N_2_. The newly exposed SiO_2_ background is then incubated for 30 minutes with 0.1 mg/mL of PAA-g-PEG-N_3_ or PAA-g-PMOXA as described above.
Incubation with PAA-g-PEG-N_3_ allows for the selective immobilization of DBCO-labelled ligands on the background for dual-ligand patterning. Alternatively, incubation with PAA-g-PMOXA allows for an anti-fouling background and single-ligand nanopatterns.

#### Preparation of the Custom-made Adhesive 96-well Plates

Bottomless 96-well plates (ID: 655000) and corresponding lids (ID: 656178) were purchased from Greiner Bio-One. Double-sided medical pressure-sensitive adhesives tested for *in vitro* cytotoxicity according to ISO 10993-5 were purchased from AdhesiveResearch, Inc. (ARcare 90106NB). This double-sided adhesive was cut to cover the 96-well plate without covering the well bottom, using a Kern Laser Systems model KER52100-i-401 HSE CO_2_-laser at 2% power with 1.2 inches/second speed (Design vector available upon request). The laser-cut adhesive was then attached to the plate, and the plate was sterilized in 70% EtOH for at least 30 minutes before removal of the last adhesive protective layer and attaching the nanopatterned coverslip.

#### Protein Labeling

Attaching the linker-tags to the proteins was done in 8-12 times molar excess of N-Hydroxy succinimide (NHS) esters with either DBCO or Biotin functional group (Sigma-Aldrich & BroadPharm). To facilitate the functional groups’ accessibility, at least a PEG4 spacer between the functional group and NHS ester was chosen. Similarly, NHS esters with fluorescent dyes (Lumiprobe – Germany) were mixed at the same time in similar molar excess to achieve labeling of the functional group and a fluorescent dye at the same time.
The reaction was either done in a PBS buffer or 50 mM HEPES with the addition of 150 mM NaCl at pH 8.0 for 1 hour at room temperature with shaking. The excess NHS linkers were removed using Amicon centrifugal filters with an appropriate molecular cut-off. Absorbance measurements determined the final protein concentration and degree of labeling on a NanoDrop® microvolume spectrophotometer.

#### Bioligand Incubation on the Nanopatterned Coverslips

Bovine serum albumin (BSA) was purchased from Sigma-Aldrich (A7030), human fibronectin was purchased from Chemicon (FC010), Streptavidin was purchased from VWR (E497), monomeric Streptavidin was purchased from Sigma-Aldrich (SAE0094), Staph. Protein A was purchased from ThermoScientific (101006), VCAM-Fc was purchased from SinoBiological (50163-m03h), Human E-Cadherin-Fc was purchased from Chemtronica (ECD-H5250)

A mixture of bioligands of interest was added to the surface in a PBS buffer for the proof-of-concept fluorescence microscopy experiments. DBCO-BSA at a final 100 μg/mL concentration and SA at 25 μg/mL. To ensure the full coverage of DBCO-BSA reaction with the N_3_ groups on the surface, the incubation was done overnight at 37°C. SA was incubated at 25 μg/mL for 30 minutes for experiments where SA was used as a linker protein. Biotinylated Protein A was incubated at 10 μg/mL for 1 hour. E-Cadherin Fc was incubated overnight at RT at 5 μg/mL. Thermally annealed 3’ Biotin functionalized 23’mer DNA to another 3’ cyanine 3’ labeled complementary DNA (Integrated DNA Technologies) was incubated at 100 nM for 30 minutes. For the small molecule patterning proof-of-concept experiments DBCO-5’FAM (Lumiprobe) and sulfo-Cyanine3-PEG3-Biotin (Lumiprobe) were incubated for 2 hours at room temperature at 1 μM.

The Protein A – monomeric Streptavidin conjugate was prepared by NHS-labelling the Protein A with methyl-tetrazine and monomeric Streptavidin with Trans-Cyclooctyne. The Amicon purified proteins were then allowed to react by the biorthogonal inverse electron-demand Diels-Alder cycloaddition and purified using a Cytiva Superdex 200 size-exclusion column on a Cytiva ÄKTA Pure protein purification system.

#### Surface Plasmon Resonance

Biacore SIA Au chips were purchased from Cytiva. The gold-covered chips were sputter-coated with 4 nm Ti, followed by 30 nm SiO_2_ to resemble the used SiO_2_/glass substrate surface chemistry. Buffers and solutions were degassed by sonication and filtered with a 0.22 μm syringe filter prior to use. A Biacore 3000 was used for the measurements. All experiments are done with a flow rate of 5 μL/min at 25°C with similar concentrations as the static conditions described above.

#### Scanning Electron Microscopy (SEM)

An FEI Magellan 400 SEM was used to acquire images based on secondary electron detection. The presented SEM images were acquired by an incoming electron beam of 5 kV and a nominal beam current of 50 pA.

#### Atomic Force Microscope (AFM)

Bruker Dimension Edge with RTESP-300 tip with a nominal tip radius of 8 nm was driven to 270-300 KHz for measurements done in intermediate mode (tapping mode™) in air. All data were plane-fitted through 3 points, and lines were aligned by modulus in Gwyddion^[2]^ before analysis or presentation.

#### X-ray Photoelectron Spectroscopy (XPS)

XPS data acquisitions were performed using a Kratos AxisUltraDLD instrument (Kratos Analytical) equipped with a monochromatic Al kα X-ray source, operating at 10 kV and 15 mA (150 W). Survey spectra (binding energy (BE) range of 0−1100 eV with a pass energy of 160 eV) were obtained to determine the presence of an element on the surface. All data presented is based on the average of 3 scans of a single spot. The acquired data were analyzed CASA XPS^[3]^.

#### Cell Culture and Proof-of-concept Experiment

C2C12 cells, obtained from the American Tissue Type Culture Collection, were cultured in high glucose DMEM (Gibco) at 37°C with 5% CO_2_ and supplemented with 10% (v/v) fetal bovine serum (FBS), 50 mg ml^−1^ penicillin, 50 mg ml^−1^ streptomycin. The cells were passaged by trypsinization when reaching approximately 80% confluence and used within passage 15 and 21. The proof-of-concept experiment was done in similar conditions by adding recombinant mouse EGF (SinoBiological, 50482-MNAY), but without FBS for 2 hours at a seeding density of 4500 cells/cm^2^. The nanopatterned 96-well palte was prepared as described above with PAA-g-PEG-Biotin and PAA-g-PEG-N_3_. A brief 0.25 mg/mL BSA blocking step was introduced to fill any potential holes on the surface before incubation with the patterned proteins. The cells were fixed for 10 minutes in 4% PFA and permeabilized in 0.2% Triton-X 100 for 10 minutes. For nucleus staining, F-Actin staining was done with 50 nM Atto-565 Phalloidin (AttoTec, Germany) and 300 nM DAPI (Sigma-Aldrich) for 30 minutes.

#### DNA-PAINT Super-Resolution Microscopy & Total Internal Reflection Fluorescence Microscopy (TIRFM)

An Oxford Nanoimager S in TIRF mode was used for DNA-PAINT^[4]–[6]^(DNA-point accumulation in nanoscale topology) super-resolution imaging. The same setup was used for TIRF imaging of DNA and protein nanopatterns. For DNA PAINT imaging, DBCO-terminated 7xR3 docking strands (IDT) were conjugated to the azide-modified Streptavidin and purified by Amicon Ultra 30 kDa Centrifugal Filter.
Cy3B-conjugated R3 imager (Metabion) was used at a concentration of 50 pM in imaging buffer C+ (PBS with the addition of 500mM NaCl & 0.05% Tween-20) with the addition of oxygen scavengers, according to the published protocol by Schnitzbauer et al.^[5]^.
561 nm laser with a power density of ∼ 2.5 kW/cm^2^ was used to illuminate the sample, and 10.000 frames with an exposure time of 100 ms were acquired. The images were drift-corrected using 90 nm Au fiducial markers and reconstructed in Picasso^[5]^.

#### Confocal Laser Scanning Microscope (CLSM)

A Zeiss LSM 700 with oil immersion (n=1.518) 64x and 100x objectives with an aperture of 1 airy unit was used to image the protein patterns or the interaction of the cells with the nanopatterned surfaces. Lasers were set to 2%, and no digital offset was applied. The images were contrast-corrected and exported by FIJI^[7]^.

Widefield Inverted Fluorescence Microscope
An inverted Olympus IX-81 with a 40x objective and a Halogen lamp was used for the proof-of-concept widefield imaging.

High-Throughput Imaging
High-throughput imaging is done with a 10x objective by an ImageXpress Pico (Molecular Devices). 50% of each well was imaged and tiled to form the two-channel fluorescence images used for the analysis. Light source intensity, exposure time, and detector sensitivity were kept constant throughout the plate imaging for fluorescence signal quantifications.

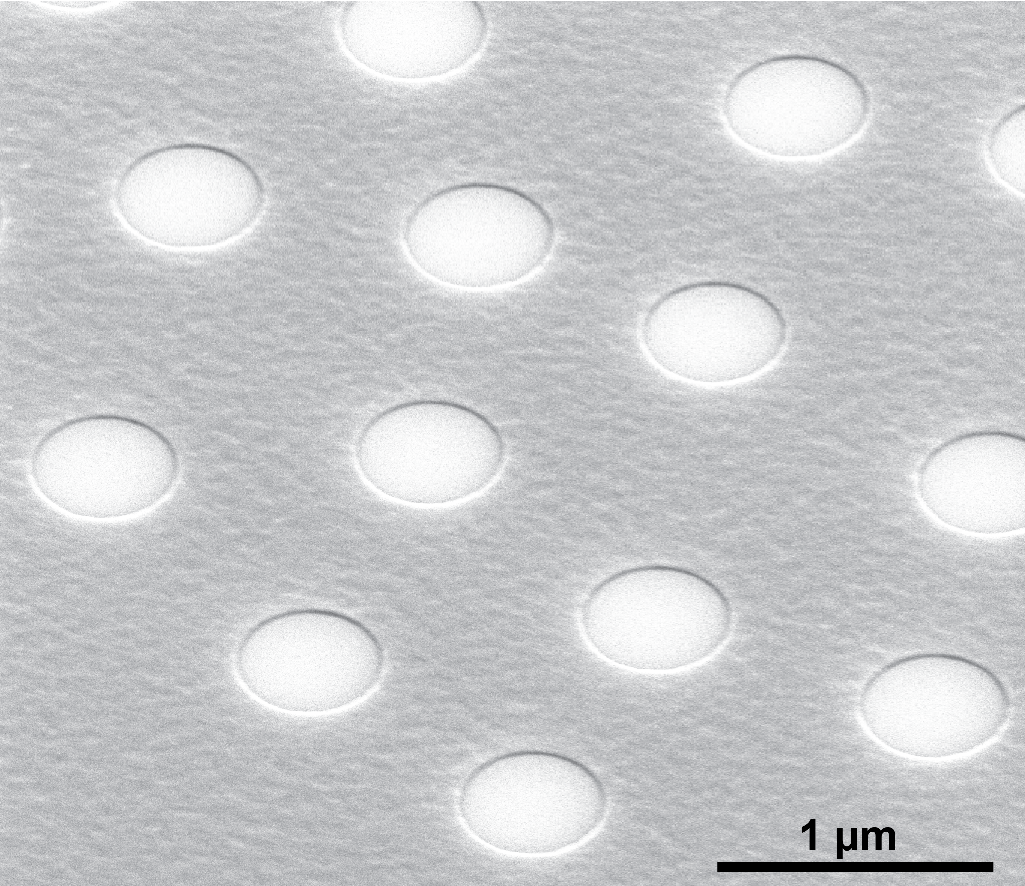

**Figure S1.** SEM sideview of the nanoparticles projection on a Si-substrate after Cr layer deposition and tape stripping the nanoparticles off the substrate. (Fabrication step C)

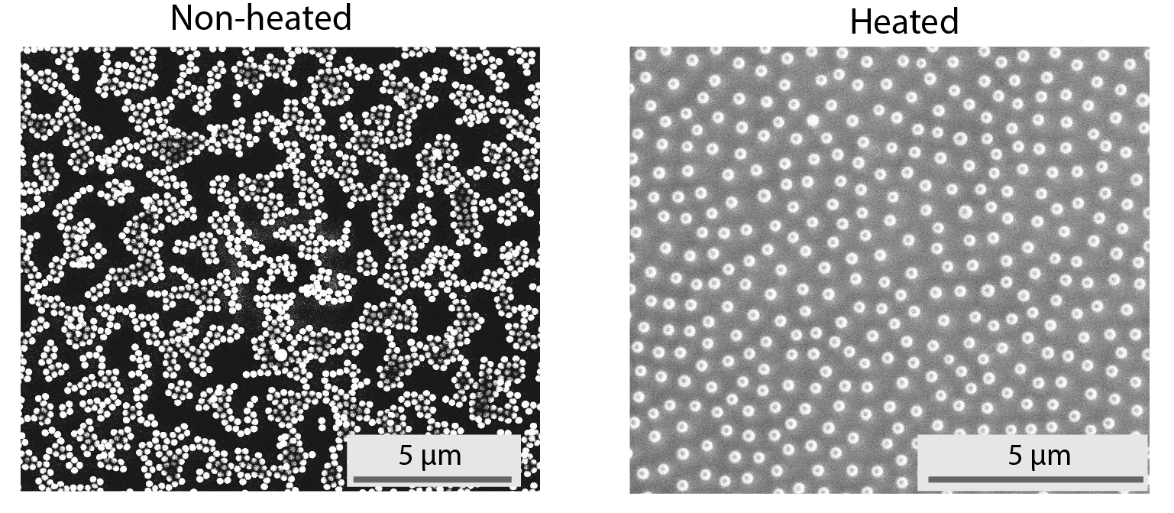

**Figure S2.** Effect of the capillary forces on adsorbed nanoparticles. Left (non-heated) shows the aggregation artifact due to the capillary forces moving the particles together. By heating the particles (Heated; right side) above their glass transition temperature, the particles adhere more strongly to the surface and can withstand the capillary forces included by drying the substrates after deposition of the nanoparticles. This effect is seen for particles that are above 200 nm in size, and there is no need to heat substrates with sub 150-200 nm particles.

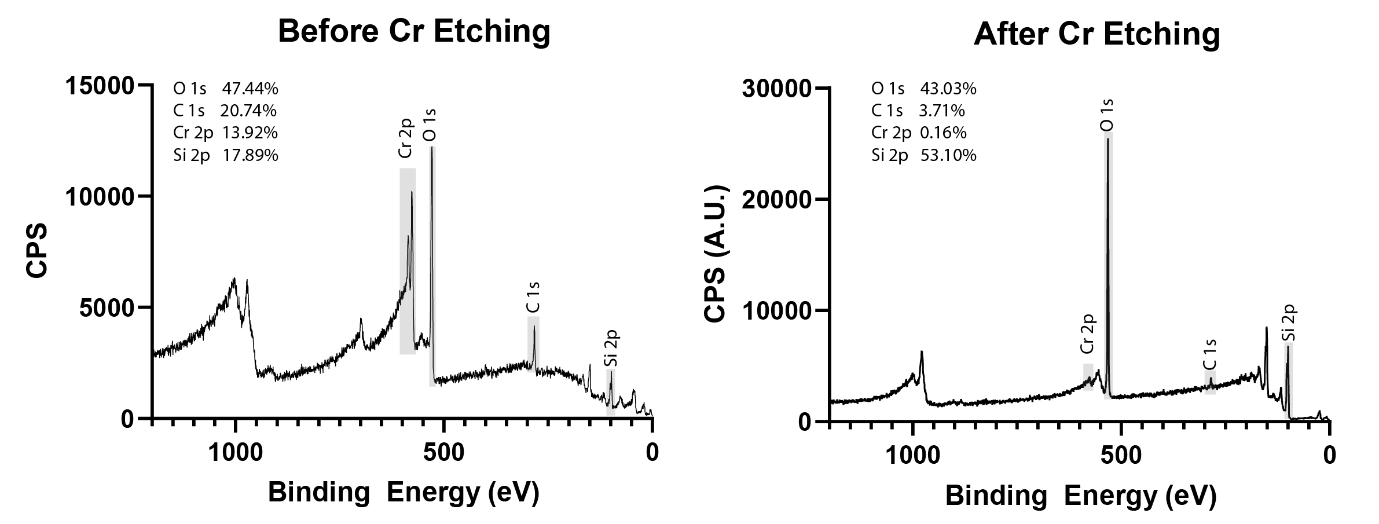

**Figure S3.** XPS surveys before and after 45 seconds Cr etching of a 200 nm patterned substrate. No PEG deposition was done for these substrates.

| a) 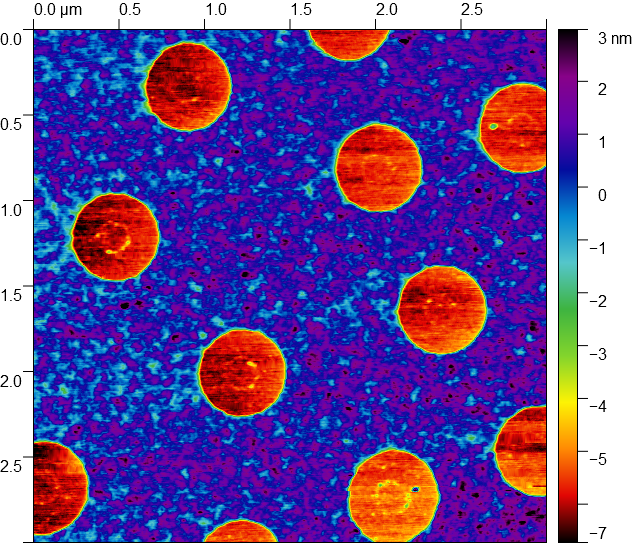 | b) 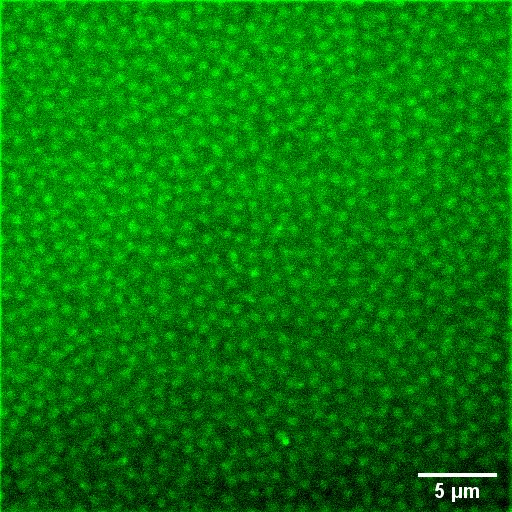 |
| --- | --- |

**Figure S4.** a) AFM scan of a substrate after 0.5 nm Cr deposition. The deposited metal layer is clearly not continuous, indicating the need for the deposition of a thicker layer to ensure complete coverage of the regions in between the particles. b) fluorescence image from the same surface after the remaining fabrication steps and incubation with Cy3-labelled Streptavidin. As seen in the AFM scan, there are “holes” in the 0.5 nm Cr thin film. This has resulted in the deposition of the PAA-g-PEG-Biotin polymer not only in the regions where the particles were placed but also in between them. This is also seen in the fluorescence image, where a significant background of Cy3-signal is detected from the regions between the nanoparticles.

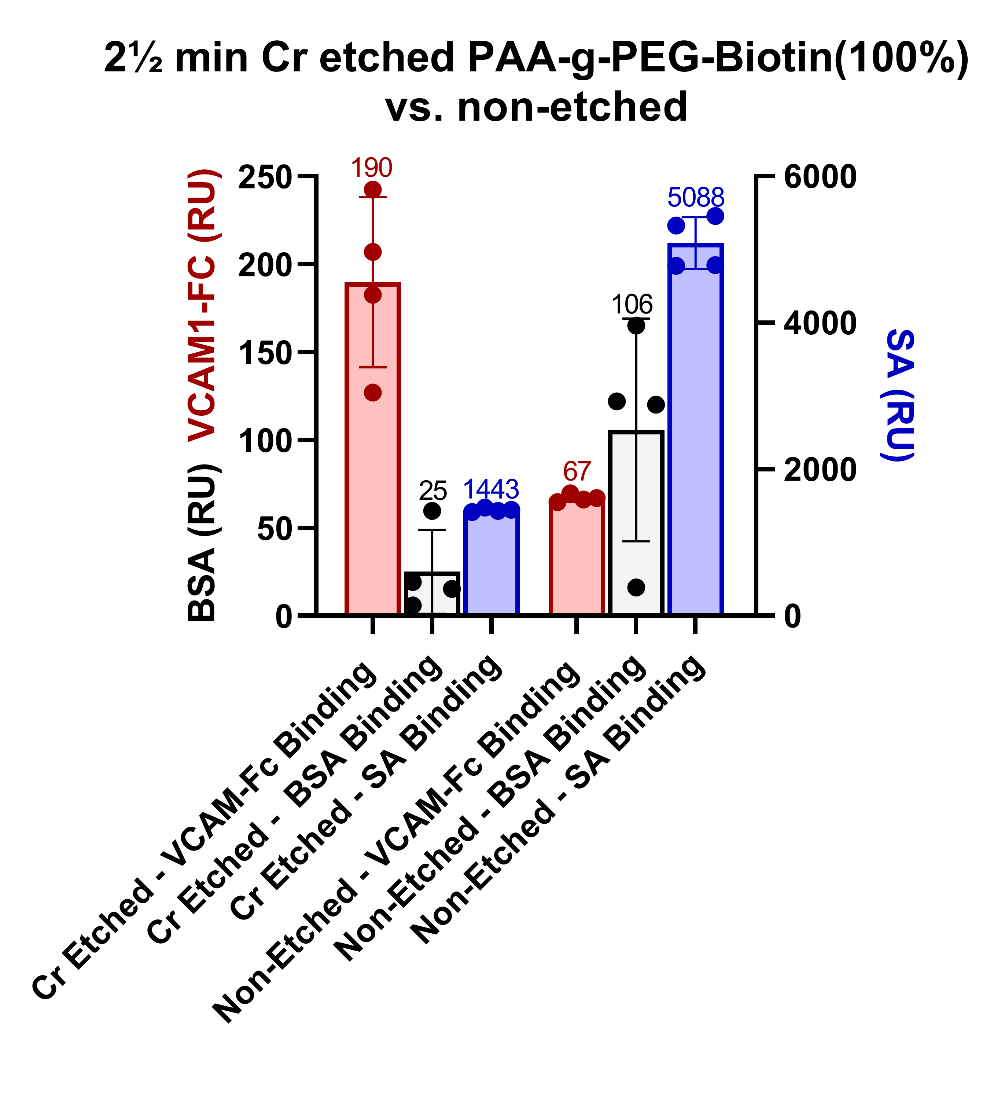

**Figure S5.** SPR results show the change in the response for 2½ min Cr etched surface compared to non-etched surface. Both surfaces are coated with PAA-g-PEG-Biotin (100%) and backfilled with PAA-g-PEG-N_3_ (100%). Gray is the amount of nonspecifically bound BSA at 0.5 mg/mL injected for 20 minutes at the flow rate of 5 μL/min. Blue is the amount of bound Streptavidin on the PEG-Biotin layer. Red is the amount of VCAM-Fc binding via Biotinylated Protein A.

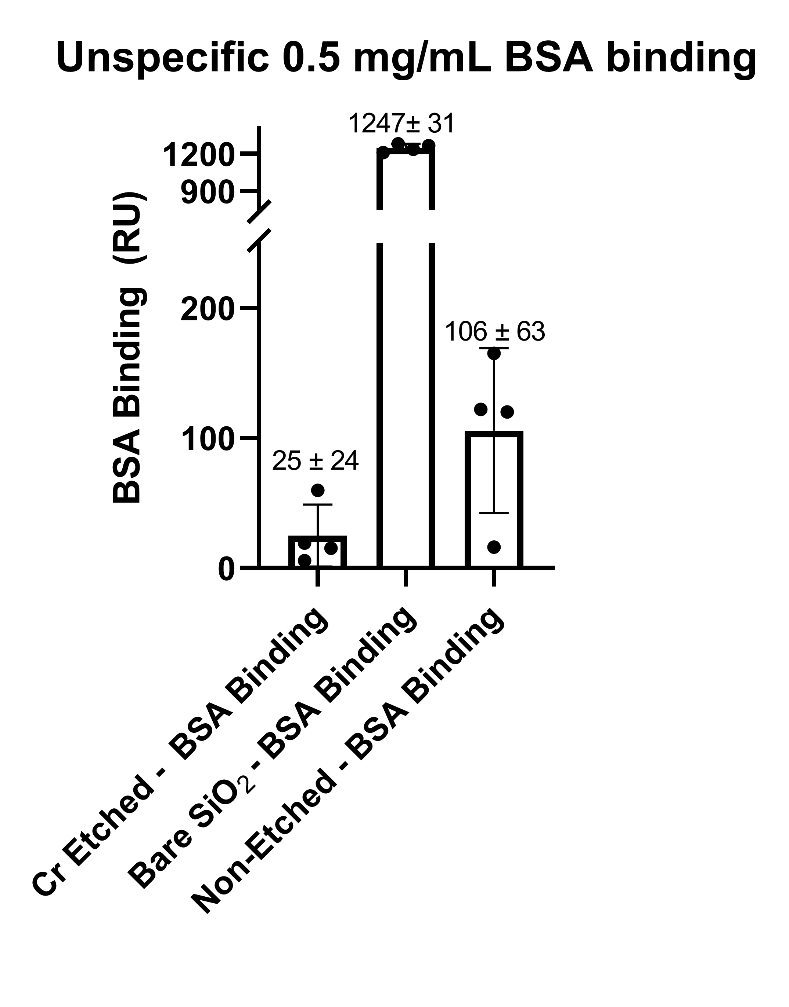

**Figure S6.** Summary of SPR sensorgram for controlling the amount of unspecifically bound BSA (0.5 mg/mL 20 min at 5 μL/min. A bare SiO_2_ chip is compared to PEGylated chips with PAA-g-PEG-Biotin (100%) and backfilled with PAA-g-PEG-N3 (100%). In (+ Etch), the surface is etched for 2½ min in Cr etchant; in (-Etch), the surface has not been exposed to the Cr etchant solution. Based on the above measurements and by normalizing the results to the bare SiO_2_ chip, anti-fouling of the etched PAA-g-PEG-Biotin surface is quantified to 98±1.7% and the non-etched surface to 91.5±4.4%.

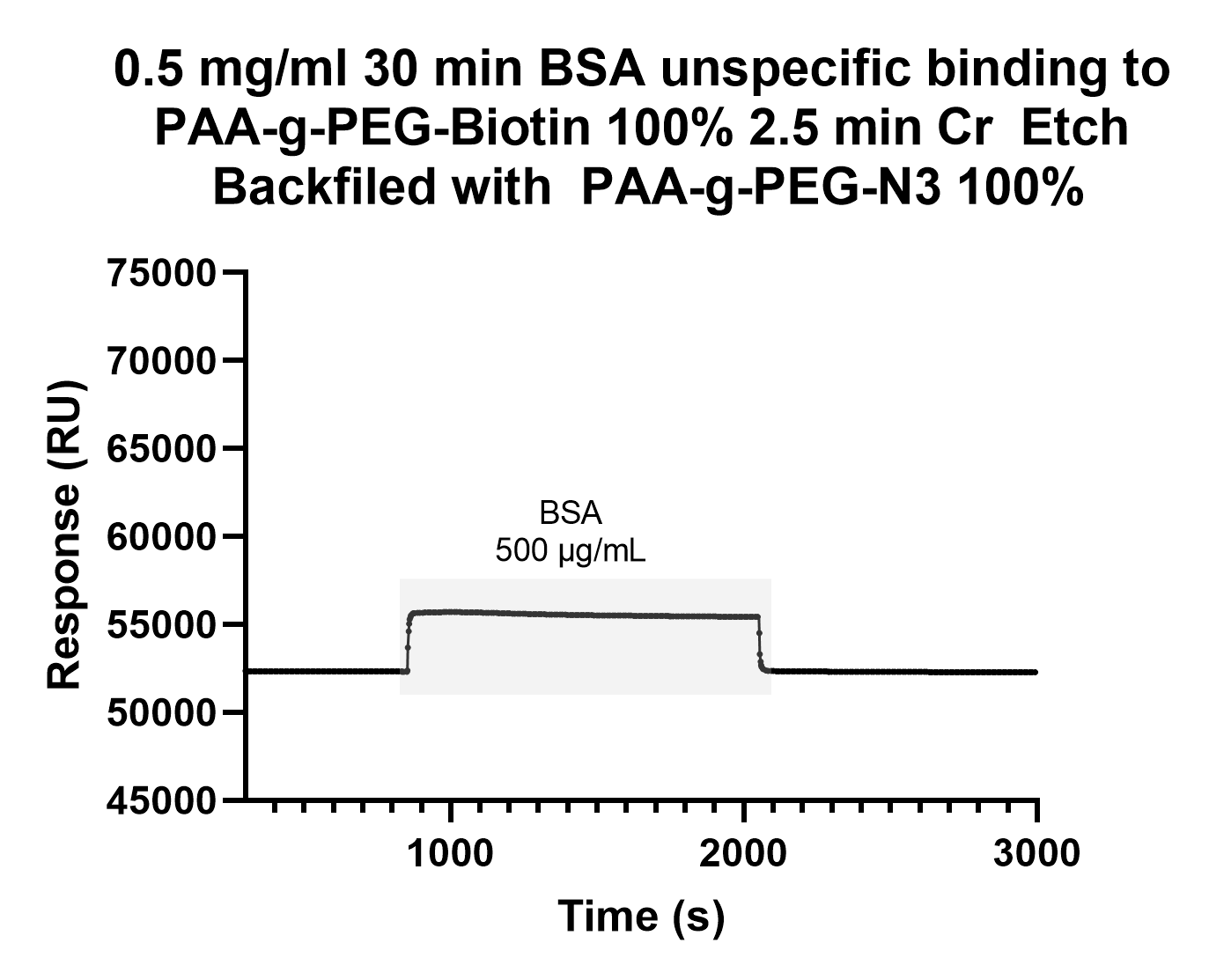

**Figure S7.** Example SPR sensorgram for the minimal unspecific binding of BSA to the etched PAA-g-PEG-Biotin surface

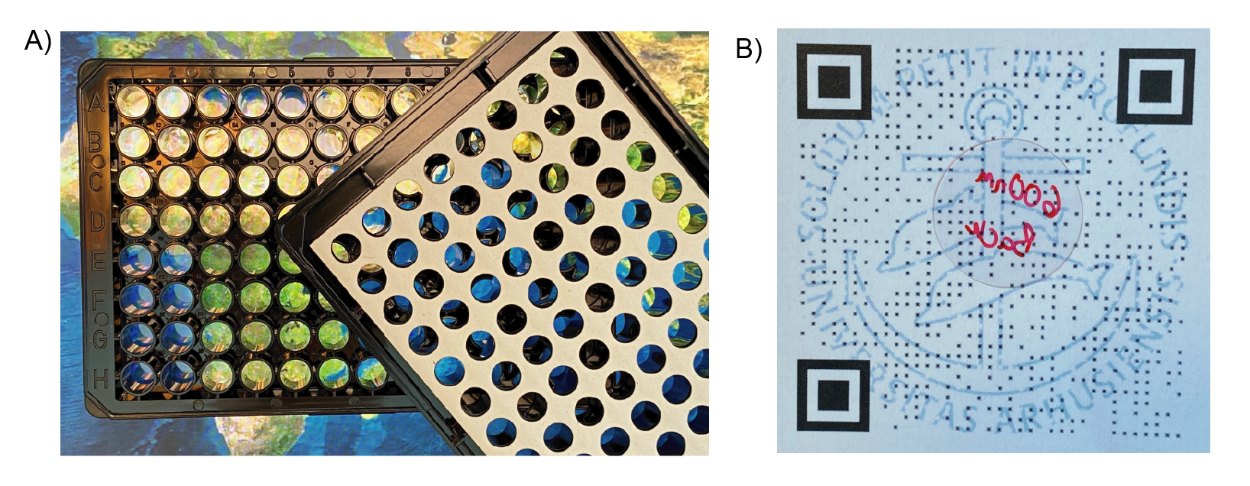

**Figure S8.** Photographs demonstrating the transparency of the assembled nanopatterned coverslip assembled to a sticky 96 well plate(a), positioned next to a sticky 96 well plate. b) Ø25 mm thin coverslip on paper with a printed QR code, still scannable.

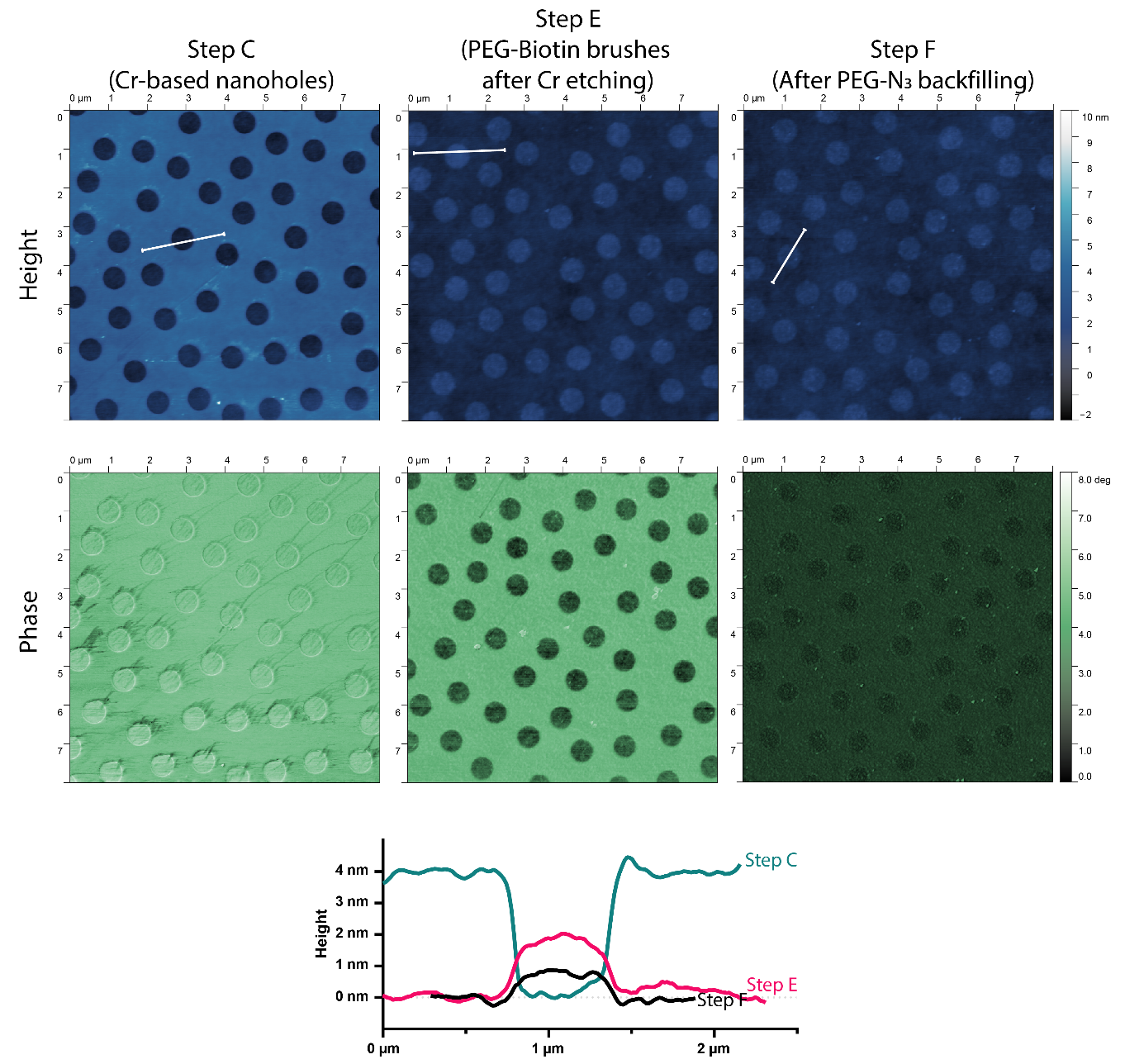

**Figure S9.** Tapping mode in air AFM scans with height and phase information for the fabrication steps C-F). After oxidation, the depth of the fabrication nanoholes is measured to approximately 4 nm. In contrast to the 0.5 nm deposited Cr layer, the deposited film is continuous without any holes in between the nanostructures. After deposition of the PAA-g-PEG-Biotin and etching away the Cr layer for 2½ min, the holes turn into bumps of approximately 2 nm. A clear change in the phase scan for these regions is seen. After backfilling the background region with PAA-g-PEG-N_3,_ the height difference for the nanostructured regions decreases from 2 nm to below 1 nm. The phase contrast is minimal in this step since the whole surface is covered with a similar polymer layer. The less than 1 nm difference in height is associated with the conjugated DBCO-PEG4-Biotin tag. The white lines indicate the position of the line profiles shown.

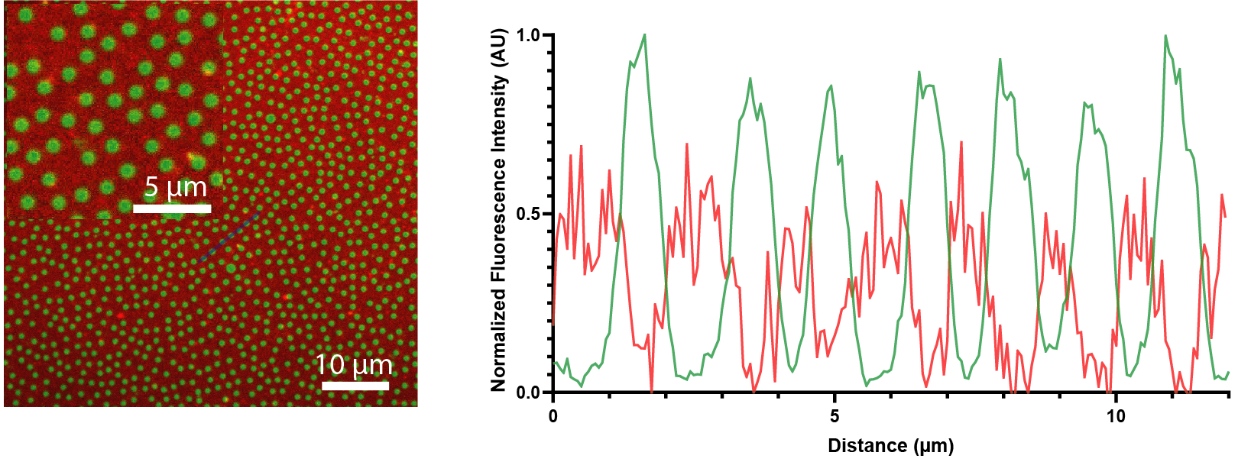

**Figure S10.** fluorescence image and line scan for the line shown in blue. There is a clear anti-correlation between where the red (DBCO-BSA-Cy5) and green (Streptavidin) signals originate from. This is quantified by Pearson correlation coefficient of -0.5, indicating a robust anti-correlation.

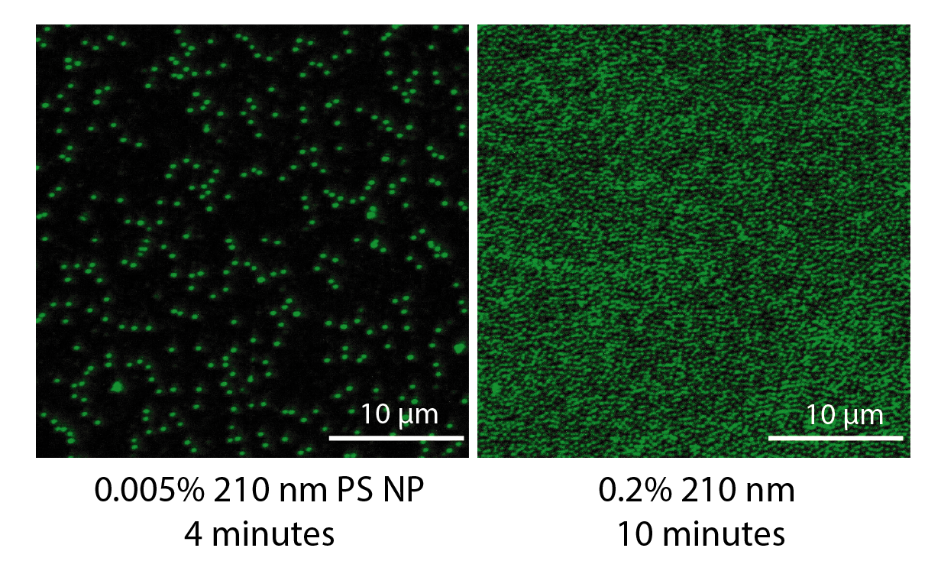

**Figure S11.** CLSM image of Streptavidin nanopatterns on 210 nm structures prepared by abruption of the particle assembly before reaching this limit (left) or allowing the particles to approach the jamming limit (right).

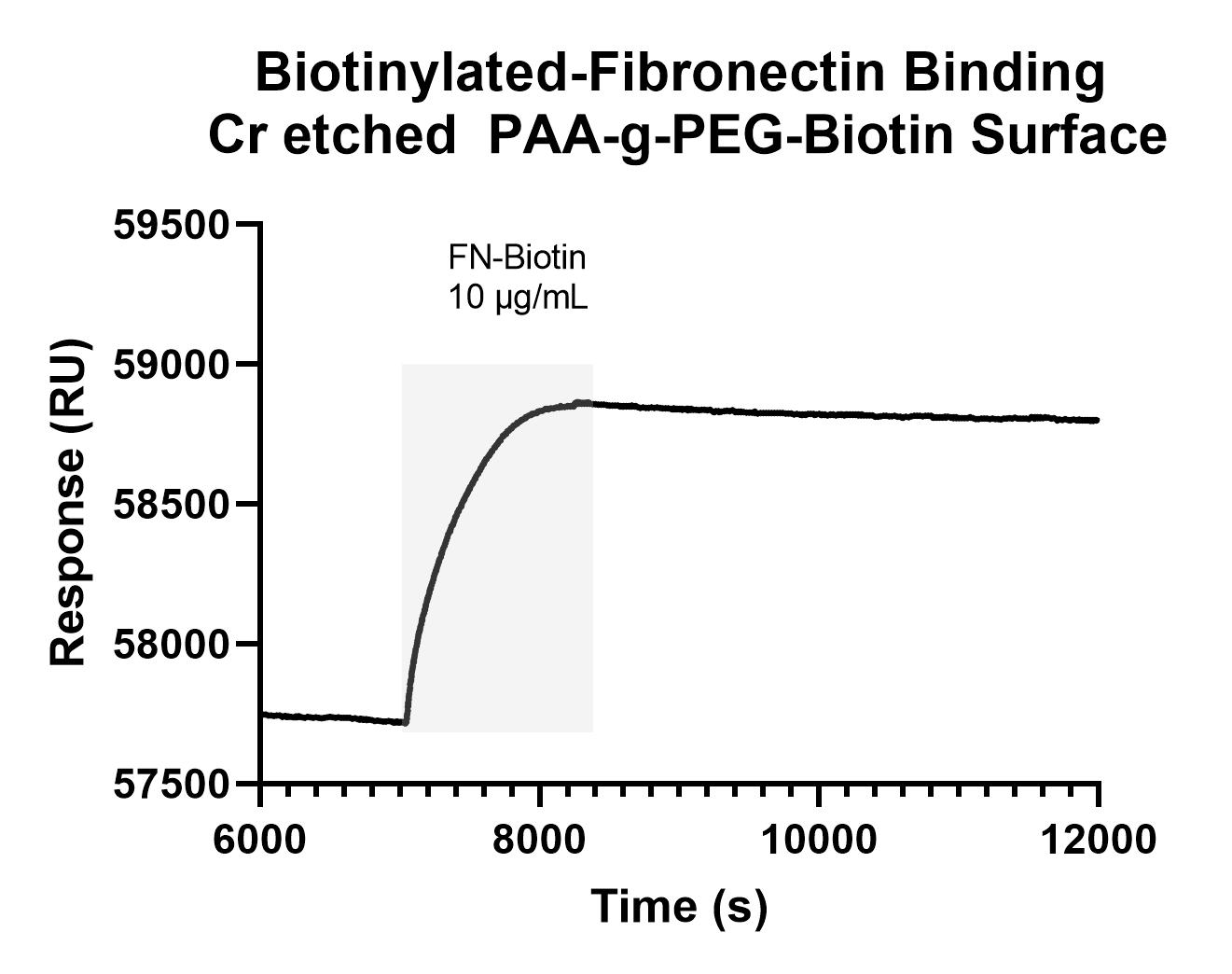

**Figure S12.** Representative SPR sensorgram from binding of Biotinylated fibronectin through Streptavidin bound to the PAA-g-PEG-Biotin brushes. Gray region indicates the injection.

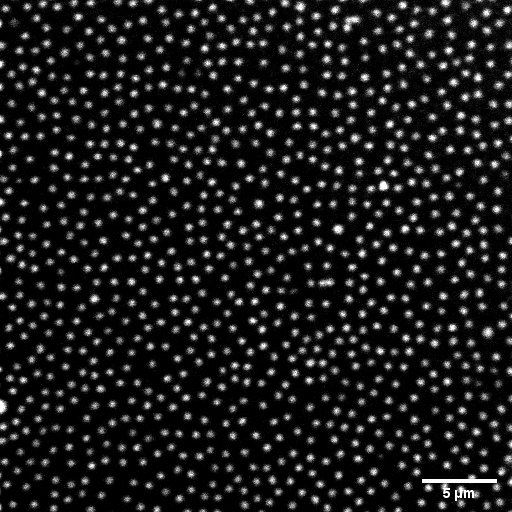

**Figure S13.** fluorescence image of Cy3-labelled Streptavidin after incubation in DMEM cell culture media with the addition of 10% serum after incubation at 37°C for 10 days.

| a) 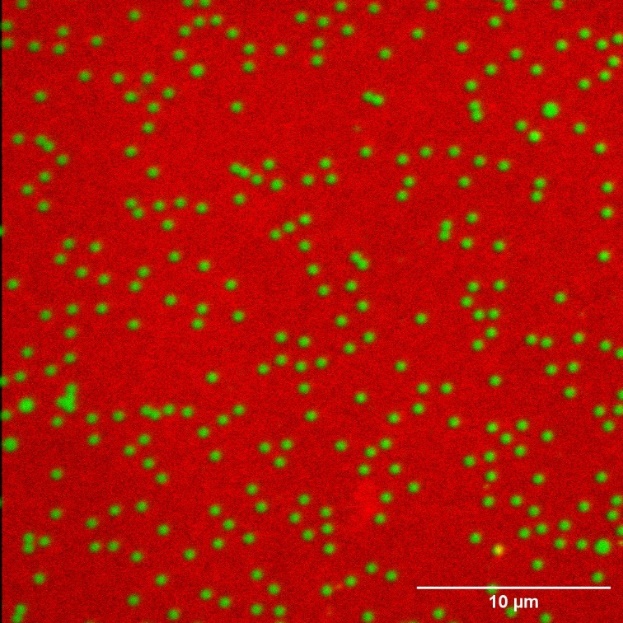 | b) 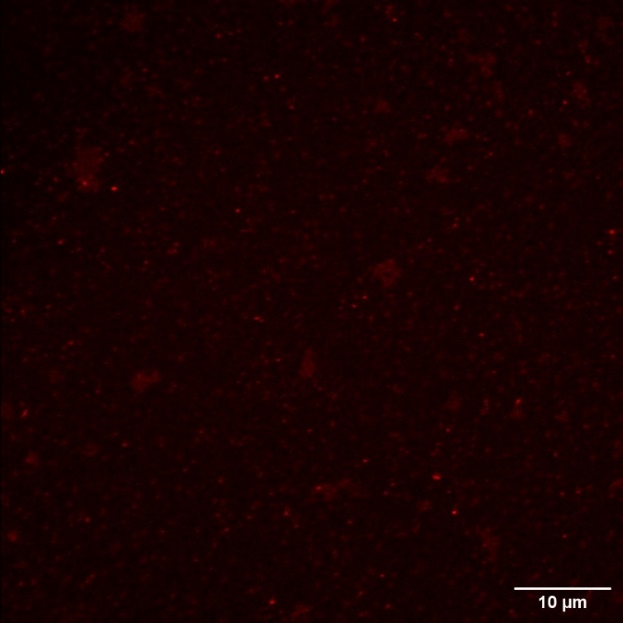 |
| --- | --- |

**Figure S14.** A 500 nm sample was prepared with PAA-g-PEG-Biotin in the nanopatterned regions and PAA-g-PEG-N_3_ in the background and stored in a -20°C freezer for 561 days before incubation with proteins. The left side figure (a) shows the proteins binding specifically to the designated regions even after 1 year of storage. Green is Cy3-labelled Streptavidin, and red is DBCO-BSA-Cy5.
On the right-side figure (b) Cy5-labelled BSA is used to control the unspecific binding. This protein was incubated overnight at 100 μg/mL. Using similar contrast as the image on the left-hand side, no signal was detectable. By increasing the laser power and detector gain, we could see a very weak signal from the surface, which mainly originated from the patterned regions with the Biotin tag.
In both cases, some aggregations of the nanostructures are visible. This is due to the aggregation of the nanoparticles in the fabrication step. Since this sample was prepared in the very early days, the particle deposition protocol was not finalized and had some aggregation and coverage issues.

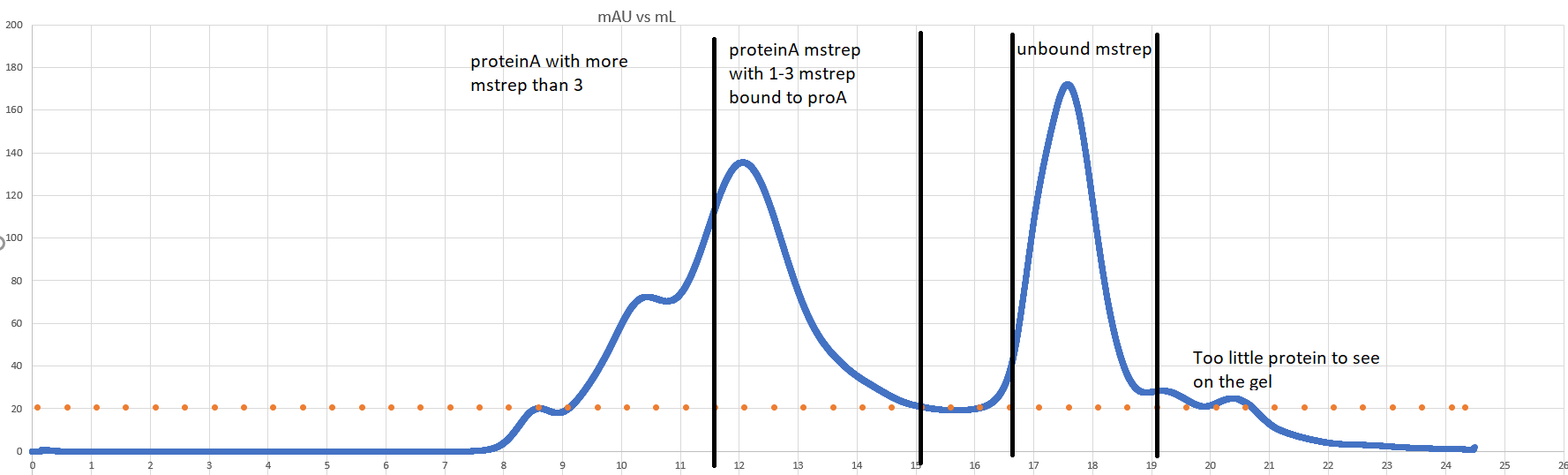

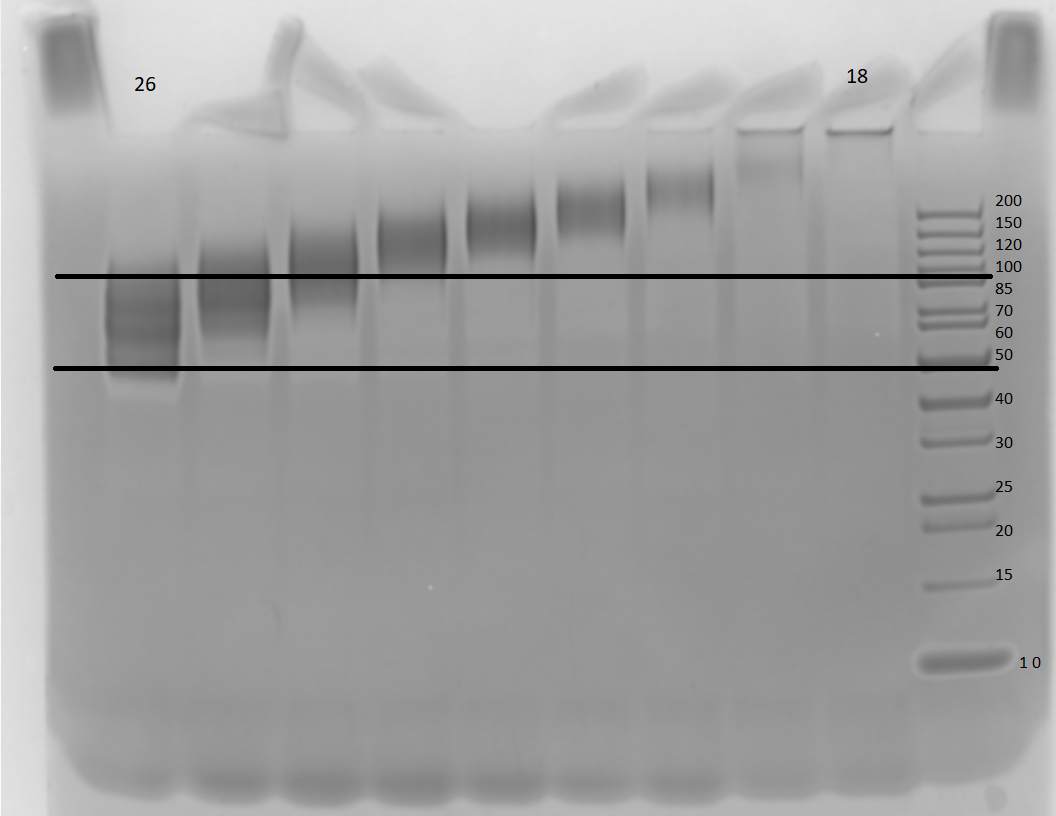

**Figure S15.** UV absorbance and the corresponding purification fractions on an SDS-PAGE gel for the Protein A-Monomeric Streptavidin conjugate purification. The purification is done by a size-exclusion column in a ÄKTA Pure Protein Purification System (Cytiva). Protein A conjugates with 1-3 Monomeric Streptavidin conjugates were collected, pooled, and used for the experiment. The purified fractions are identified as yellow dots on the chromatogram. SDS-PAGE gel displays fractions 18 to 26.

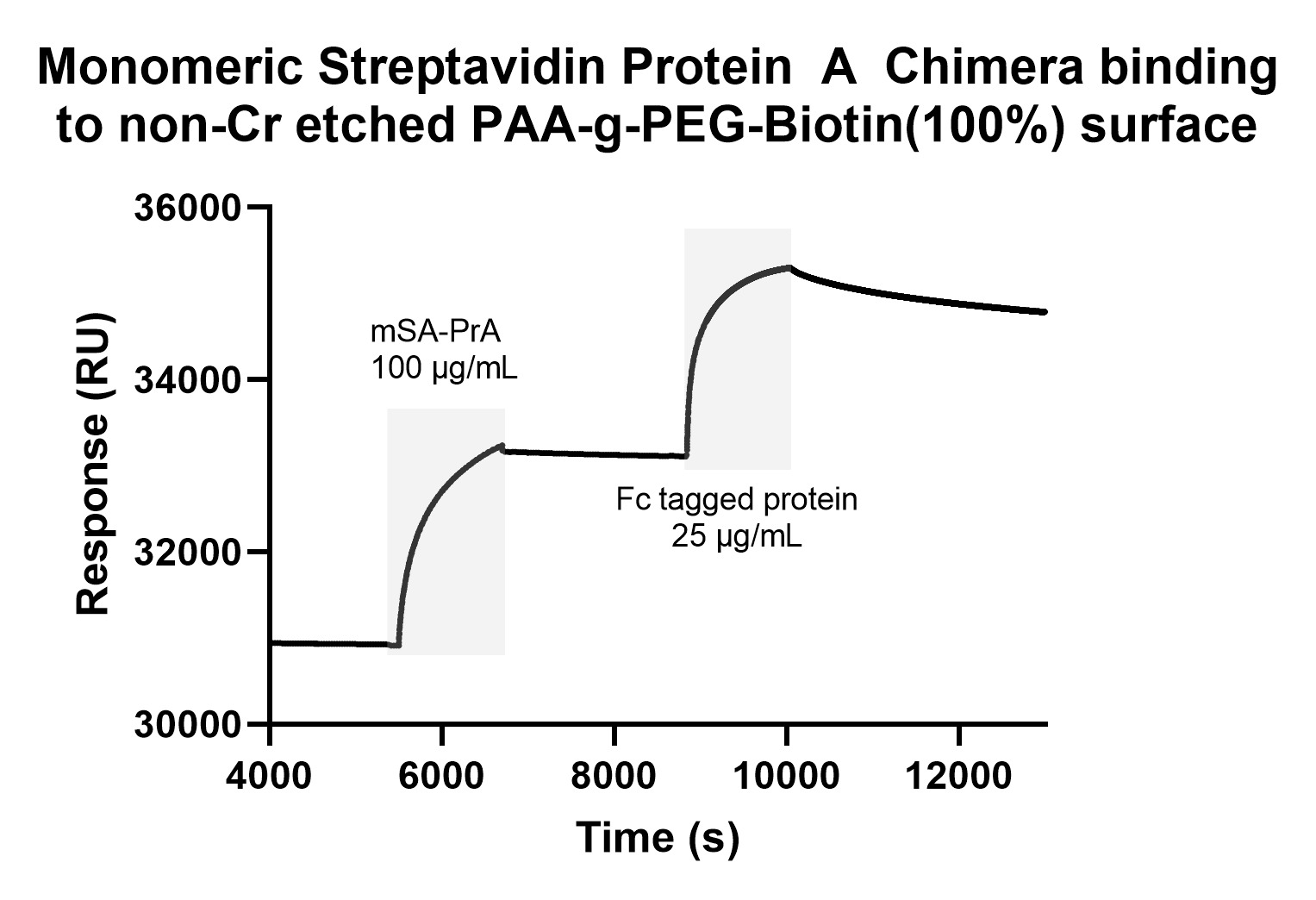

**Figure S16.** SPR sensorgram for binding monomeric Streptavidin-Protein A conjugated (mSA-PrA) to PAA-g-PEG-Biotin 100% non-Cr-etched surfaces. There is no need for the “linker” Avidin molecule since between 1 and 3 monomeric Streptavidin molecules are already covalently bound to the Protein A. The binding of a Fc tagged protein confirms the functionality of the mSA-PrA chimera.

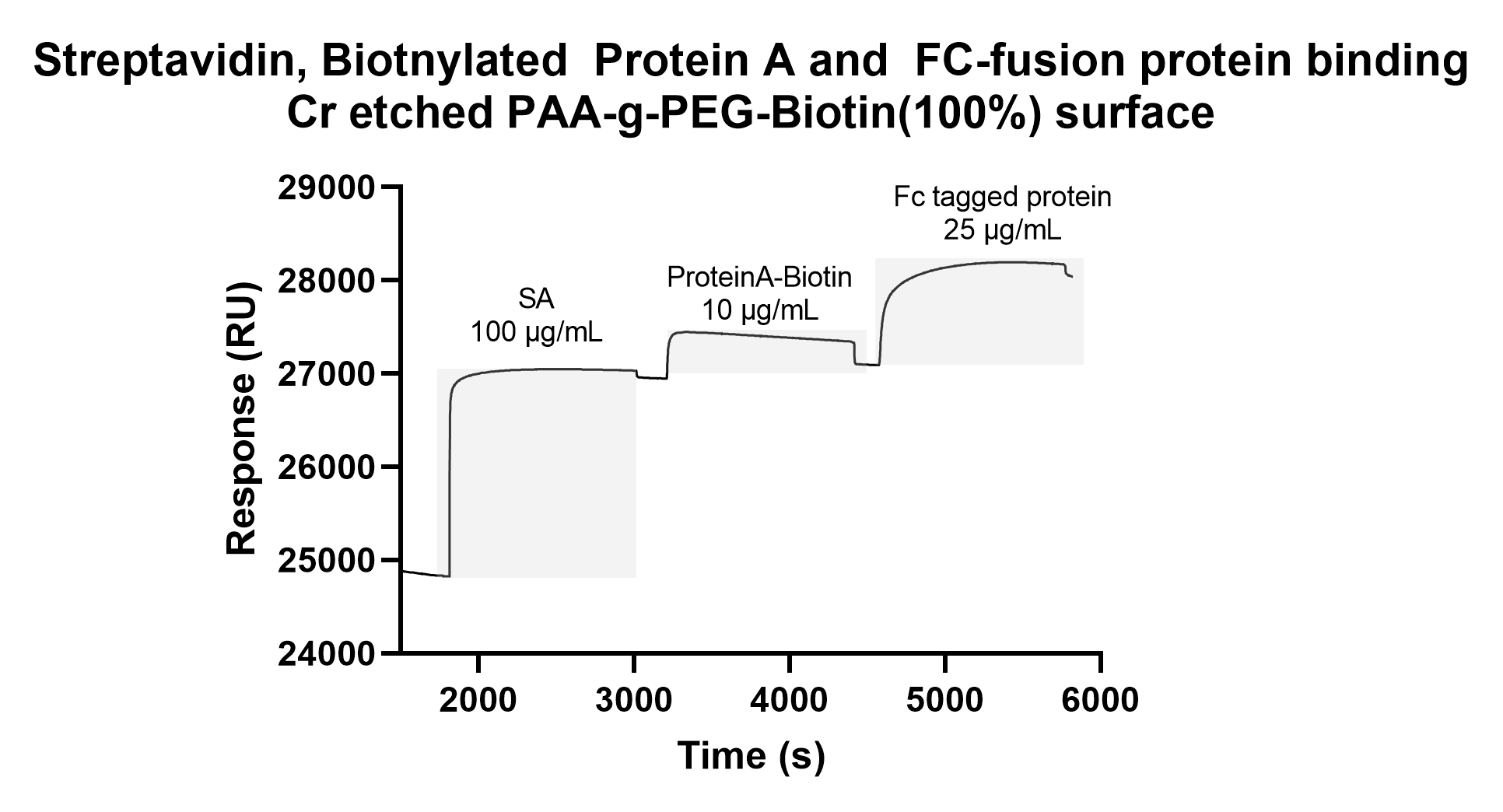

**Figure S17.** Example sensorgram for sequential and specific binding of Streptavidin (SA), Biotinylated Protein A (ProteinA-Biotin), and an Fc tagged protein. The experiment is done on a SiO_2_ SPR chip replicating the patterned region of the substrates.

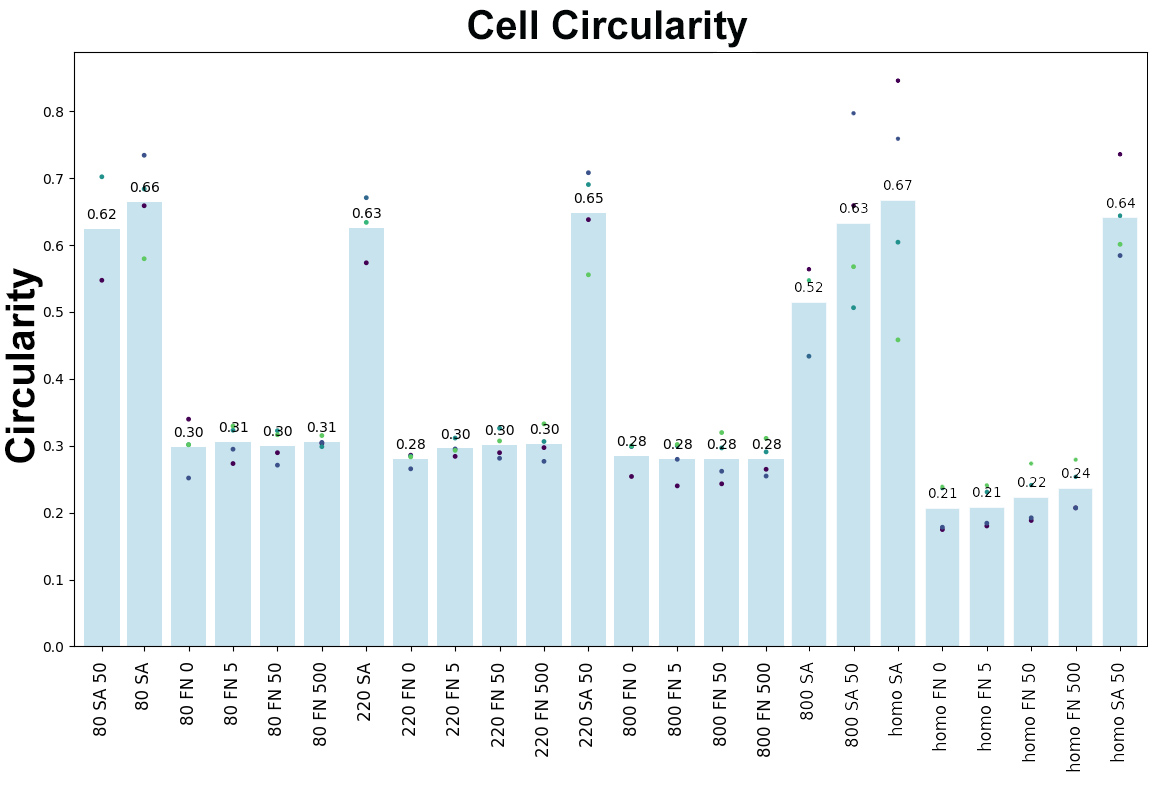

**Figure S18.** Cell Circularity – each dot represents a well for each condition. The negative controls without the adhesive protein on Streptavidin (SA) with or without EGF are much more circular than the cells on positive controls (FN). This suggests that the few cells (less than 2%) that are bound on the negative controls cannot spread and thus are circular. A slight increase in circularity is also seen when EGF concentration is increased on the nanopatterned fibronectin substrates.

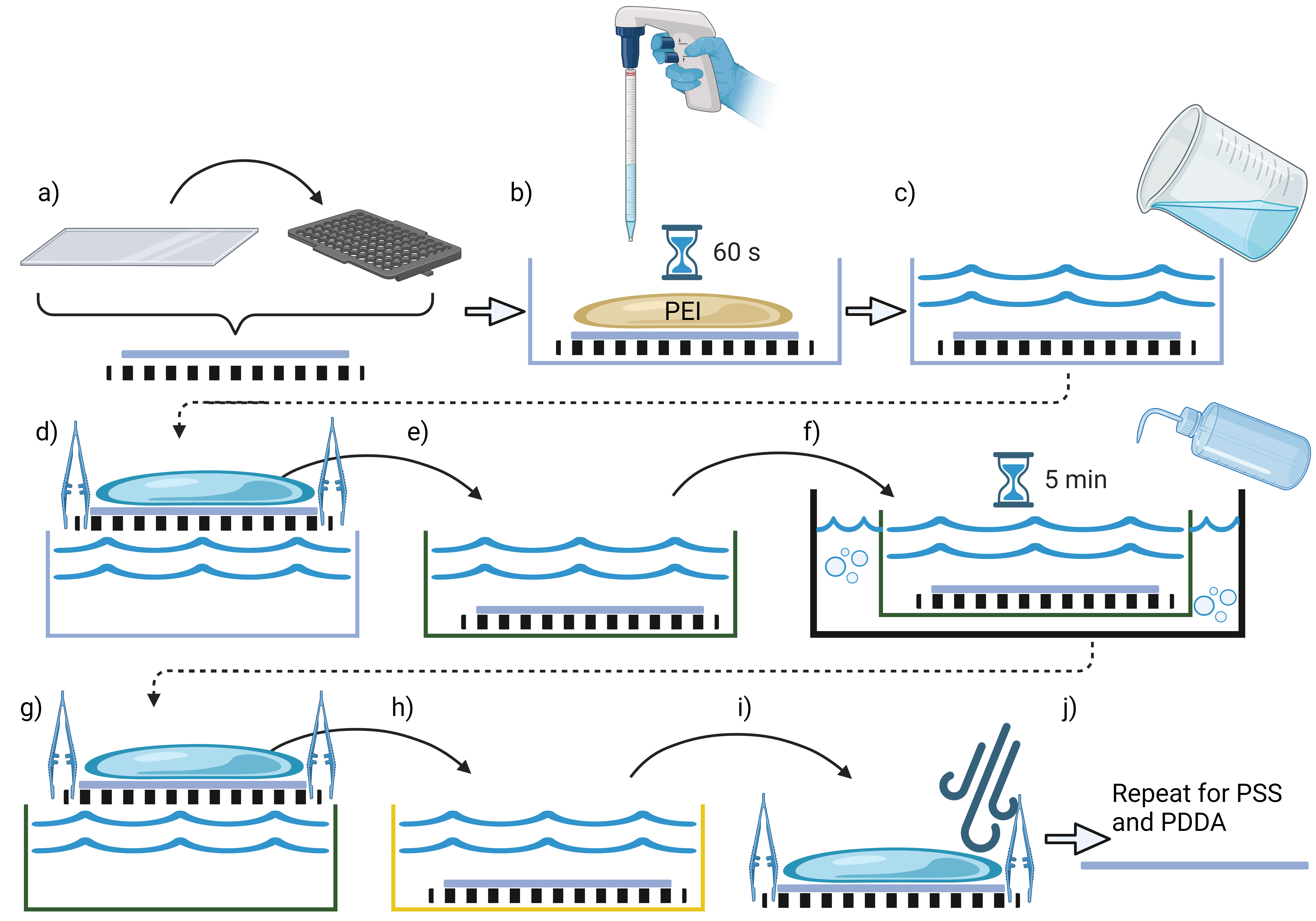

**Figure S19.** Flowchart displays polyelectrolytes (PEs) deposition on large glass substrates. a) First, the glass is cleaned by ultrasonication in acetone, rinsed in isopropanol, and placed on a sample holder made from a rack of pipette tips with embedded metallic weights to avoid floating. b) The sample on the holder is put into a glass container, and the first PE is added with a large pipette in one continuous motion. c) DI H_2_O is poured on top of the sample in one continuous motion until the sample is fully submerged. d-e) The sample is then transferred into another container filled with DI H_2_O, keeping a droplet on the sample to avoid dewetting. f) The container is now put in an ultrasonic bath and sonicated for 5 minutes while spraying the sample gently with DI H_2_O a few times during the sonication. g-h) The sample is transferred into a new container with clean DI H_2_O. i) The sample is transferred onto a table covered with paper towels and dried with an N_2_ gun. The drying is done using a 3D-printed nozzle adapter, which covers one side of the sample and makes it possible to dry the whole surface in one continuous motion. This procedure is now repeated for the remaining PSS and PDDA PE layers.

 
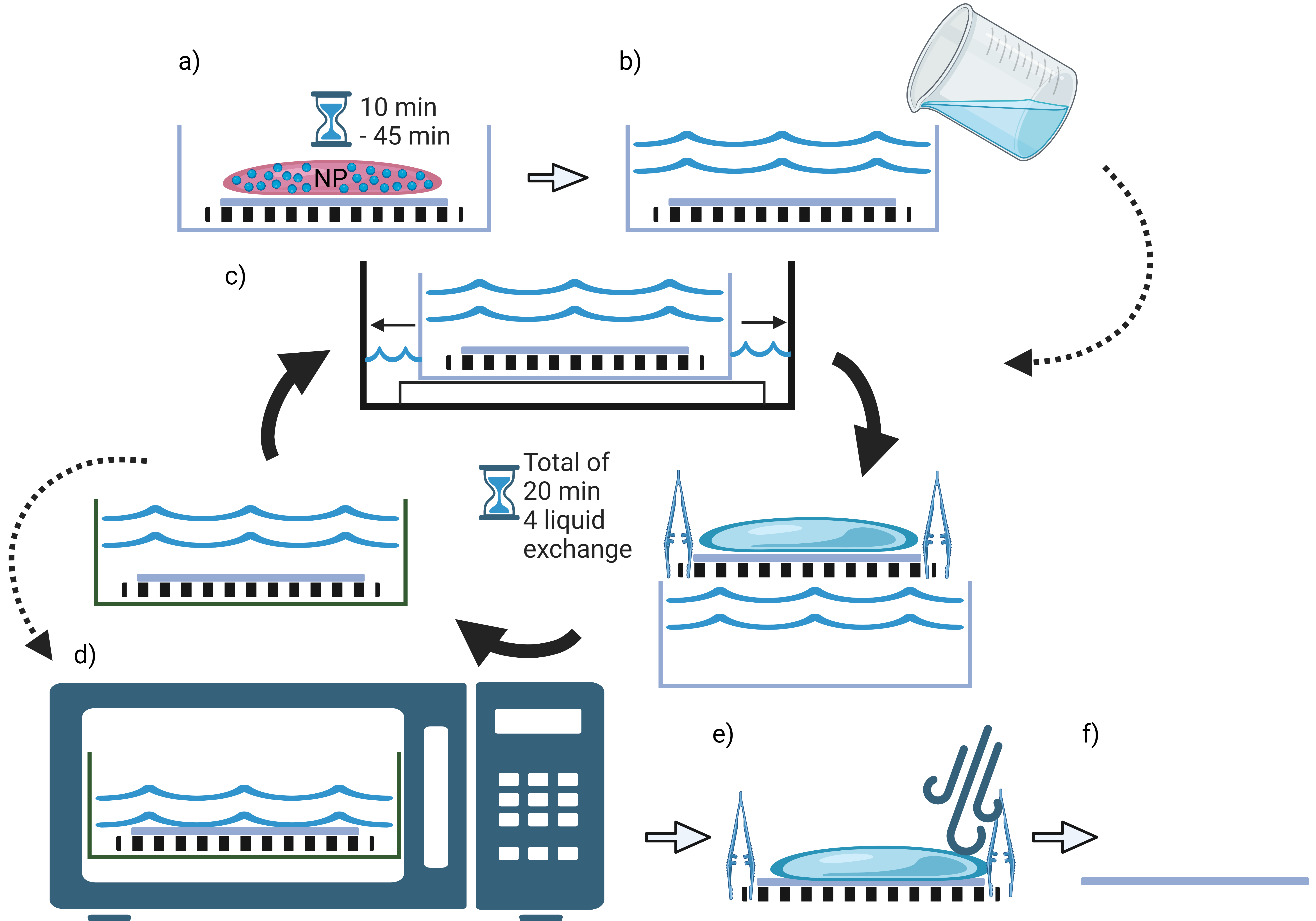

**Figure S20.** Flowchart displays nanoparticles' deposition onto large glass samples, which are covered with the polyelectrolyte layers described in S19. a) First, the sample already covered with polyelectrolytes is put onto a sample holder in a glass container, and the nanoparticle solution is added in one continuous motion. b) The solution (0.1 % nanoparticles) is left to incubate for 10-45 minutes depending on the particle size (10 min for 100 nm, 20 min for 300 nm, 30 min for 600 nm, and 45 minutes for 800 nm). Leftover particles are washed off by gently pouring DI H_2_O on the sample until fully submerged. c) This is followed by shaking in a water bath (Julabo SW23) set to shake at 70 rpm. d) During the 20-minute shake, the liquid is exchanged four times. Hereafter, the particles were heated in a microwave oven at 900 W for 10 minutes. e) Lastly, the sample was put onto a table covered with paper towels and dried with N_2_ gun using the 3D-printed nitrogen nozzle adapter. f) This results in the finished sample, ready for Cr mask deposition.
